# *Vicia faba* TPC1, a genetically encoded variant of the vacuole Two Pore Channel 1, is hyperexcitable

**DOI:** 10.1101/2021.12.22.473873

**Authors:** Jinping Lu, Ingo Dreyer, Miles Sasha Dickinson, Sabine Panzer, Dawid Jaslan, Carlos Navarro-Retamal, Dietmar Geiger, Ulrich Terpitz, Dirk Becker, Robert M. Stroud, Irene Marten, Rainer Hedrich

## Abstract

To fire action-potential-like electrical signals, the vacuole membrane requires the depolarization-activated two-pore channel TPC1, also called <u>S</u>lowly activating <u>V</u>acuolar SV channel. The TPC1/SV channel, encoded by the *TPC1* gene, functions as a voltage-dependent and Ca^2+^-regulated potassium channel. TPC1 currents are activated by a rise in cytoplasmic Ca^2+^ but inhibited by luminal Ca^2+^. In search for species-dependent functional TPC1 channel variants, we studied polymorphic amino acids contributing to luminal Ca^2+^ sensitivity. We found that the acidic residues E457, E605 and D606 of the Ca^2+^-sensitive Arabidopsis AtTPC1 channel were neutralized by either asparagine or alanine in *Vicia faba* and many other Fabaceae as well. When expressed in the Arabidopsis loss-of-AtTPC1 function background, the wild type VfTPC1 was hypersensitive to vacuole depolarization and insensitive to blocking luminal Ca^2+^. When AtTPC1 was mutated for the three VfTPC1-homologous polymorphic site residues, the Arabidopsis At-VfTPC1 channel mutant gained VfTPC1-like voltage and luminal Ca^2+^ insensitivity that together made vacuoles hyperexcitable. These findings indicate that natural TPC1 channel variants in plant families exist which differ in vacuole excitability and very likely respond to changes in environmental settings of their ecological niche.

**Significance statement:** Vacuolar electrical excitability and stress-related Ca^2+^ signaling depends on the activity of the vacuolar cation channel TPC1. Until now, the regulatory features of AtTPC1 from the model plant *Arabidopsis thaliana* was believed to apply to the TPC1 channels of other species. However, here we now show that, surprisingly, the VfTPC1 channel of the economic broad bean, in contrast to AtTPC1, proves to be hyperactive and confers hyperexcitability to the vacuole. The different gating behavior is most likely related to an impaired Ca^2+^ sensor site in the vacuolar pore vestibule, rising the probability to open at more negative membrane voltages. These natural variants of the TPC1 channel could help the plant adapt and respond to environmental challenges.

## Introduction

Soon after the patch-clamp technique was first applied to plant cells (for review see 1, 2), the <u>s</u>low <u>v</u>acuolar SV channel was identified as a calcium-regulated, voltage-dependent cation channel (3). After two further decades it was demonstrated that the *Arabidopsis thaliana* SV channel is encoded by the single copy gene *TPC1* (4) and that TPC1 contributes to long-distance electrical and Ca^2+^ signaling and is an essential player in vacuole excitability (5–7).

Within the tree of live, there are TPC1-like sequences in animals and plants (see for review 8, 9). Remarkably, during land plant evolution characteristic structural fingerprints of TPC1-type channels remained unchanged from mosses to flowering plants (10–12). Recently, both the crystal and cryoEM structure of the Arabidopsis AtTPC1 channel and thus the molecular topology of this vacuolar cation channel became available (13–16). AtTPC1 is formed by a dimer whose monomers consist of two tandem Shaker-like cassettes, each with six transmembrane domains (TM1-6), connected by a cytoplasmic loop with two EF hands. Calcium binding to the EF hands is necessary for SV/TPC1 channel activation at elevated cytosolic Ca^2+^ levels (15, 17). Structural 3D motif comparison and point mutation analysis identified the major voltage-sensing domain (VSD) required for depolarization-dependent activation in each monomer in the first four transmembrane segments of the second Shaker-like cassette (13, 18). On a similar experimental basis, it was found that Ca^2+^ binding to the EF hands opens the channel gate at elevated cytosolic Ca^2+^ levels (13, 17). CryoEM structures further suggested that activation of VSD2 is required for Ca^2+^ activation of the EF hand domain (15). In contrast, channel activation is strongly suppressed when the Ca^2+^ level in the vacuolar lumen reaches 1 mM and more (5, 13). Luminal Ca^2+^ can bind to three non-canonical calcium binding sites each formed by three acidic residues in AtTPC1 (site 1: D240, D454, E528; site 2: E239, D240, E457; site 3: E605, D606 and D607; 13, 14, 16). Site 1/2 residues are located in luminal linker regions between transmembrane domains while site 3 residues are found in the luminal pore entrance. Among them, site 1 and 3 play key roles in voltage gating and luminal Ca^2+^ inhibition.

In a screening with chemically induced Arabidopsis mutants for plants that produce elevated amounts of the wound hormone jasmonate (JA), the TPC1 mutant *fou2* (fatty acid oxygenation upregulated 2) was identified (19). Interestingly, the *fou2* channel behaves like a hyperactive TPC1 channel (19, 20) and opens already close to the vacuole resting voltage, whereas wild type AtTPC1 becomes active only upon depolarization. The hypersensitivity of the *fou2* TPC1 channel towards voltage results from a mutation of the negatively charged glutamate at position 454 to the uncharged asparagine (D454N). This TPC1 site is directed towards the vacuole lumen and is part of the non-canonical Ca^2+^ sensor site 1 (12–14). Therefore, elevated luminal Ca^2+^ levels inhibit the activity of wild type TPC1 channels but not of the *fou2* mutant channel (21). Consequently, *fou2* vacuoles must experience episodes in which the membrane potential is short-circuited. As a result, *fou2* plants appear wounded and produces large amounts of jasmonate (19). Furthermore, they accumulate an increased amount of Ca^2+^ in the vacuole (20). Functional studies in the context of the *fou2* mutation site showed that additional luminal amino acids other than the central Ca^2+^ binding sites like Asp454 are involved in luminal Ca^2+^ sensing, e.g. the adjacent E457 from Ca^2+^ sensor site 2 (12–14). Neutralization of E457 desensitized AtTPC1 towards vacuolar Ca^2+^ but unlike the *fou2* mutation did not additionally promote channel opening (12). The TPC1 channels from *Vicia faba* and *Arabidopsis thaliana* are historically established prototypes of the SV channel. Nevertheless, only AtTPC1 but not VfTPC1 has been identified at the molecular level so far. To fil this gap, we cloned VfTPC1 and analyzed its electrical properties. Despite similarities of the fava bean channel with its Arabidopsis homolog, both TPC1s showed also astonishing differences in Ca^2+^-dependent properties. In search for the structural reasons for this divergence, we could pinpoint three polymorphic sites within two luminal Ca^2+^ coordinating regions. When these polymorphic sites were implemented into the Arabidopsis TPC1 channel, AtTPC1 was converted into a hyperactive SV/TPC1 channel desensitized to luminal Ca^2+^, mimicking the properties of VfTPC1 and the AtTPC1 mutant *fou2*.

## Results

### VfTPC1 is a native hyperactive TPC1 channel variant with low luminal Ca^2+^ sensitivity

In our search for natural TPC1 channel variants, we noticed that SV currents have been recorded from *Vicia faba* guard cell vacuoles even under unnatural high luminal Ca^2+^ loads (50 mM; 22). Considering that hardly any AtTPC1/SV currents were recorded in *Arabidopsis thaliana* mesophyll vacuoles even at a lower luminal Ca^2+^ level such as 10 mM (21), the TPC1 channel variants of *Vicia faba* and *Arabidopsis thaliana* seem to differ in their Ca^2+^ sensitivity. In contrast to *TPC1* from *Arabidopsis thaliana* (4), the *VfTPC1* gene has not yet been identified. To gain insights into the molecular structure and function of VfTPC1, we isolated RNA from fava bean. Following RNA-seq and *de novo* transcriptome assembly, we identified a single *VfTPC1* transcript, just as in *Arabidopsis thaliana* (4). After VfTPC1 was cloned and fused to an eYFP tag, mesophyll protoplasts isolated from the Arabidopsis TPC1-loss-of-function mutant *tpc1-2* were transiently transformed with the fava bean TPC1 channel. Similar to TPC1 from Arabidopsis (Fig. S1) and other plant species (23), VfTPC1 was found to localize exclusively to the vacuole membrane as visualized by the fluorescent eYFP signal (Fig. S1). In whole-vacuole patch-clamp experiments with such fluorescent mesophyll vacuoles, macroscopic outward-rectifying SV/TPC1-like currents were elicited upon depolarizing voltage pulses (Fig. 1A). However, compared to AtTPC1, VfTPC1 channels differed in kinetics, voltage dependence and luminal calcium sensitivity (Figs. 1, 2, S2 and S3).

**Fig. 1.**
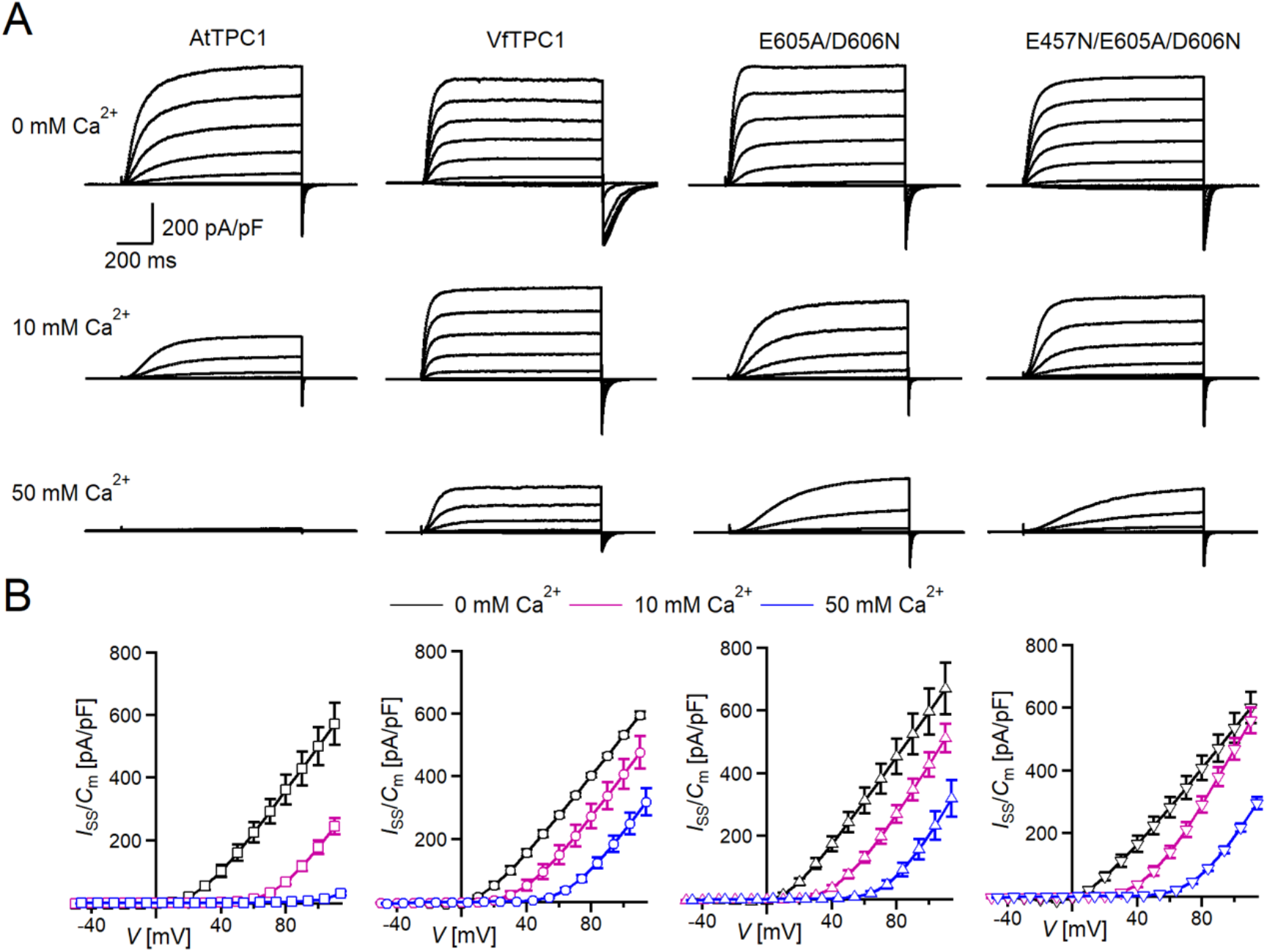
Effect of voltage and luminal Ca^2+^ on TPC1/SV currents of *Vicia faba* and *Arabidopsis thaliana* channel variants. (*A*) Macroscopic TPC1/SV current recordings from mesophyll vacuoles liberated from *Arabidopsis thaliana* protoplasts isolated from the TPC1-loss-of-function mutant *tpc1-2* and transformed with different TPC1 channel types. E605A/D606N and E457N/E605A/D606N represent AtTPC1 channel mutants. AtTPC1 and VfTPC1 denote wild type TPC1 channels from *Arabidopsis thaliana* and *Vicia faba, respectively*. TPC1/SV currents elicited upon depolarizing voltages pulses in the range −80 mV to +110 mV in 20 mV increments at indicated luminal Ca^2+^ concentrations are shown. (*B*) Normalized TPC1/SV currents (*I*_ss_/*C*_m_) derived from current recordings under different luminal Ca^2+^ conditions as those shown in *A* were plotted against the clamped membrane voltage (*V*). Symbols represent means ± SE. Squares = AtTPC1 wild type with n_0/10Ca_ = 5, n_50Ca_ = 4; circles = VfTPC1 wild type with n_0Ca_ = 5, n_10/50Ca_ = 6; upright triangles = AtTPC1-E605A/D606N with n_0/10Ca_ = 5, n_50Ca_ = 4; reversed triangles = AtTPC1-E457N/E605A/D606N with n_0/10Ca_ = 5, n_50Ca_ = 4. In *B* AtTPC1 wild-type data at 0 and 10 mM Ca^2+^ are identical to those shown in (16).

**Fig. 2.**
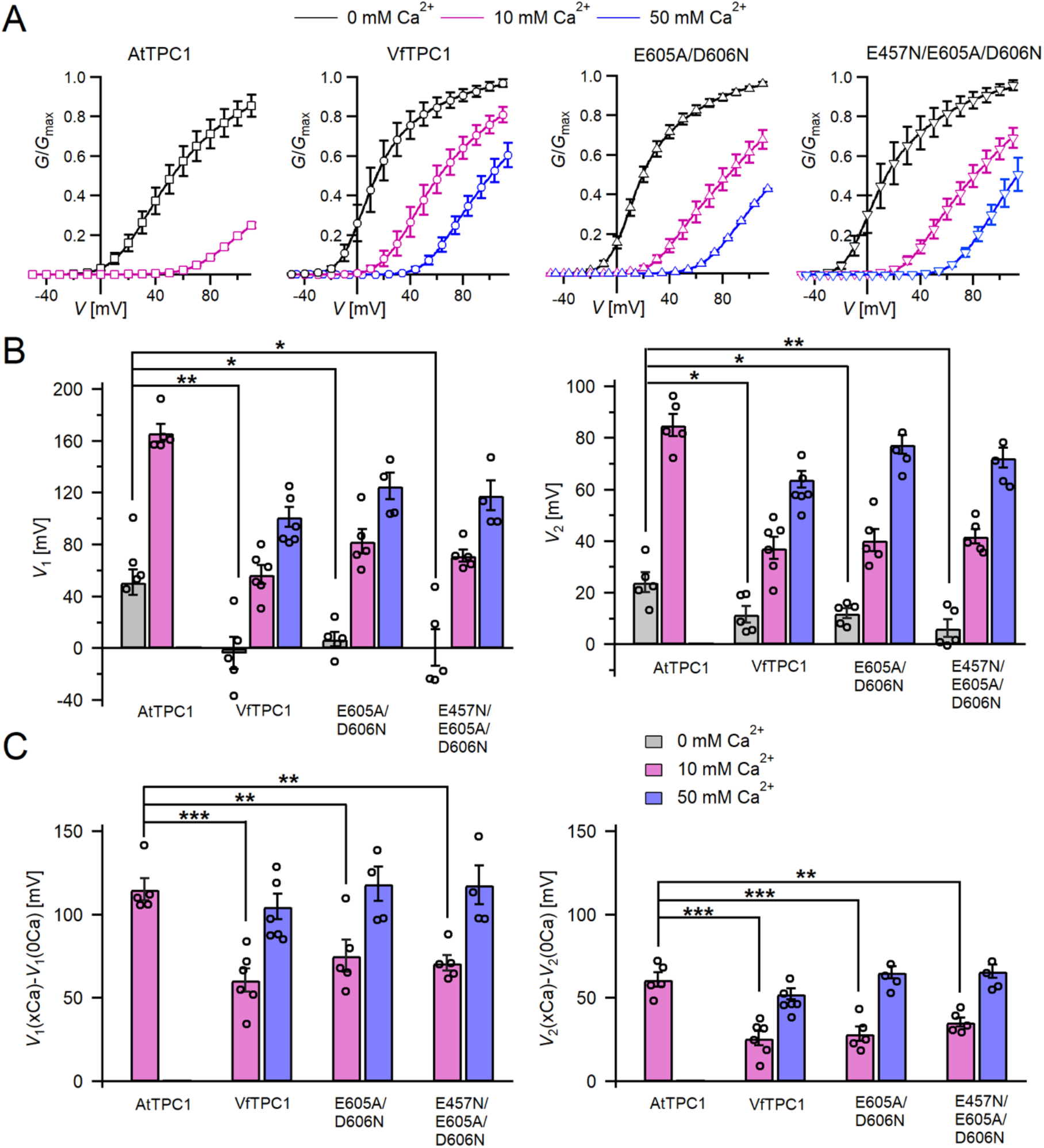
Channel activity of *Vicia faba* and *Arabidopsis thaliana* TPC1 channel variants in response to membrane voltage and luminal Ca^2+^. (*A*) conductance-voltage plots (*G*/*G*_max_ (*V*)) determined for the different TPC1 channel variants as a measure for their relative open-channel probability under indicated luminal Ca^2+^ conditions. Best fits of the *G*/*V* plots to a double Boltzmann function are given by the solid lines. Squares = AtTPC1 wild type, circles = VfTPC1 wild type, upright triangles = AtTPC1-E605A/D606N, reversed triangles = AtTPC1-E457N/E605A/D606N. (*B*) The midpoint voltages *V*_1_ (left) and *V*_2_(right) derived from the fits of the *G*/*V* plots shown in *A* are given for the different channel variants at the indicated Ca^2+^ condition. To test for significant differences between the V_1/2_ values under 0 mM luminal Ca^2+^, a statistical analysis was performed with one-way ANOVA combined with a Dunnett’s post hoc comparison test (\**p* < 0.05, \*\**p* < 0.01). (*C*) The differences in the midpoint voltages *V*_1_ (left) and *V*_2_(right) shown in (**b**) between 0 and 10 and if available between 0 and 50 luminal Ca^2+^ are shown. The changes in V_1/2_ values related to a rise from 0 to at 10 Ca^2+^ were statistically analyzed with one-way ANOVA together with a Dunnett’s post hoc comparison test (\*\**p* < 0.01; \*\*\**p* < 0.001). In *B* and *C* individual data points were inserted as open black circles into the bar chart. In *A-C* the number of experiments (n) was as follows: AtTPC1 wild type n_0/10Ca_ = 5; VfTPC1 wild type with n_0Ca_ = 5, n_10/50Ca_ = 6; AtTPC1-E605A/D606N n_0/10Ca_ = 5, n_50Ca_ = 4; AtTPC1-E457N/E605A/D606N n_0/10Ca_ = 5, n_50Ca_ = 4. Data in *A-C* represent means ± SE. AtTPC1 wild-type data at 0 and 10 mM Ca^2+^ are identical to those shown in (16).

In the absence of luminal Ca^2+^, the current activation was faster and the current deactivation slower with VfTPC1 than with AtTPC1 (Figs. 1A, S2A and B, S3A and B). A closer look at the current-voltage curves further revealed that in addition to outward currents, inward currents were also triggered by voltages in the range of −40 and 0 mV, but only from vacuoles harboring VfTPC1 and not AtTPC1 (Fig. S4A). This points to a shift in the voltage activation threshold for VfTPC1 channel opening by about 30 mV to more negative voltages. For further quantification of this effect, the voltage-dependent relative open-channel probability curves (*G/G*_max_(*V*)) were determined for both TPC1 channel variants (Fig. 2A). They were fitted with a double Boltzmann equation describing the voltage-dependent channel transitions between two closed and one open state (C_2_ ⇆ C_1_ ⇆ O) (18, 24). From these fits the midpoint voltages *V*_2_ and *V*_1_ were derived (Fig. 2B), showing in particular, a significant difference in midpoint voltage *V*_1_ of VfTPC1 (−3.5 ± 12.3 mV, n=5) and AtTPC1 (51.2 ± 9.6 mV, n=5). These results confirm that VfTPC1, unlike AtTPC1, activates near the vacuolar resting membrane voltage (Fig. 2B).

When the luminal Ca^2+^ concentration was increased from 0 to 10 mM Ca^2+^, the activation kinetics of AtTPC1 was strongly slowed down. However, this effect was much less pronounced for VfTPC1. At +90 mV, for example, the half-activation time increased by about 170% for AtTPC1, but only 53% for VfTPC1 (Fig. S2A-C, E and F). Even more impressive was the fact that AtTPC1 currents were suppressed by about 50% at 10 mM luminal Ca^2+^ and completely vanished at 50 mM luminal Ca^2+^ (Fig. 1). In contrast, the VfTPC1 current densities at zero and 10 mM luminal Ca^2+^ did not differ in magnitude, and at 50 mM Ca^2+^ the VfTPC1 current density still reached about 50% of the level measured under luminal Ca^2+^-free conditions. The strongly reduced susceptibility of VfTPC1 currents to inhibitory luminal Ca^2+^ ions was associated with a significantly reduced inhibitory effect on voltage activation (Fig. 2). Compared to 0 mM Ca^2+^, at 10 mM Ca^2+^ the *V*_1_ and *V*_2_ values increased by about 115 mV and 61 mV for AtTPC1, but only by about 61 mV and 26 mV for VfTPC1, respectively (Fig. 2B and C). A rise from 10 to 50 mM luminal Ca^2+^ caused a further positive-going shift of the VfTPC1 activation voltage curve (*G*/*G*_max_(*V*)), indicated by a 1.7- and 1.6-fold rise in *V*_1_ and *V*^2^, respectively (Fig. 2B). AtTPC1 currents, however, were so small at 50 mM luminal Ca^2+^ (Fig. 1) that activation voltage curves (*G*/*G*_max_(*V*)) could not be resolved reliably. Together these findings document that VfTPC1 is much less susceptible to inhibitory luminal Ca^2+^ than AtTPC1.

### Fabaceae and Brassicaceae TPC1 channels are polymorphic in luminal Ca^2+^ sensing motifs

To gain insights into the functional domains of these Brassicaceae and Fabaceae TPC1 channel proteins underlying their different response to membrane voltage and luminal Ca^2+^, we aligned not only the AtTPC1 and VfTPC1 but also other TPC1-like amino acid sequences of these plant families. Focusing on charged residues at the three sites that are involved in luminal Ca^2+^ coordination and sensitivity in AtTPC1 (Fig. 3; (12, 14, 16), we found that these sites were highly conserved among the Brassicaceae TPC1s. Only two of the 12 TPC1 variants tested (BrTPC1, LeTPC1) exhibit a neutralizing substitution at sites homologous to the AtTPC1 pore mouth residue Asp606. In contrast, most TPC1-types of the Fabaceae species show one up to three non-conservative variations of the residues. Eleven of the 14 Fabaceae TPC1 channels harbor a non-charged asparagine or glutamine instead of the negatively charged glutamate at sites homologous to AtTPC1-E456 or −E457 (Fig. 3). Remarkably, this E456Q/E457N polymorphism was additionally accompanied by the neutralization of one or two negatively charged residues at site 3 within the luminal pore mouth in all Fabaceae TPC1 channels, with the exception of *Bauhinia tomentosa* BtTPC1 (Fig. 3). In this regard, the Fabaceae TPC1 channels group in two clusters, containing either one or two non-charged residues (Ala/Val, Asn) at the homologous sites to AtTPC1-E605 and −D606 (Fig. S5). The fava bean TPC1 channel belongs to a triple polymorphism subgroup, containing alanine and asparagine at sites homologous to the Ca^2+^ sensor sites 2 and 3 of AtTPC1 (Fig. 3; AtTPC1-E457/E605/D606; VfTPC1-N458/A607/N608).

**Fig. 3.**
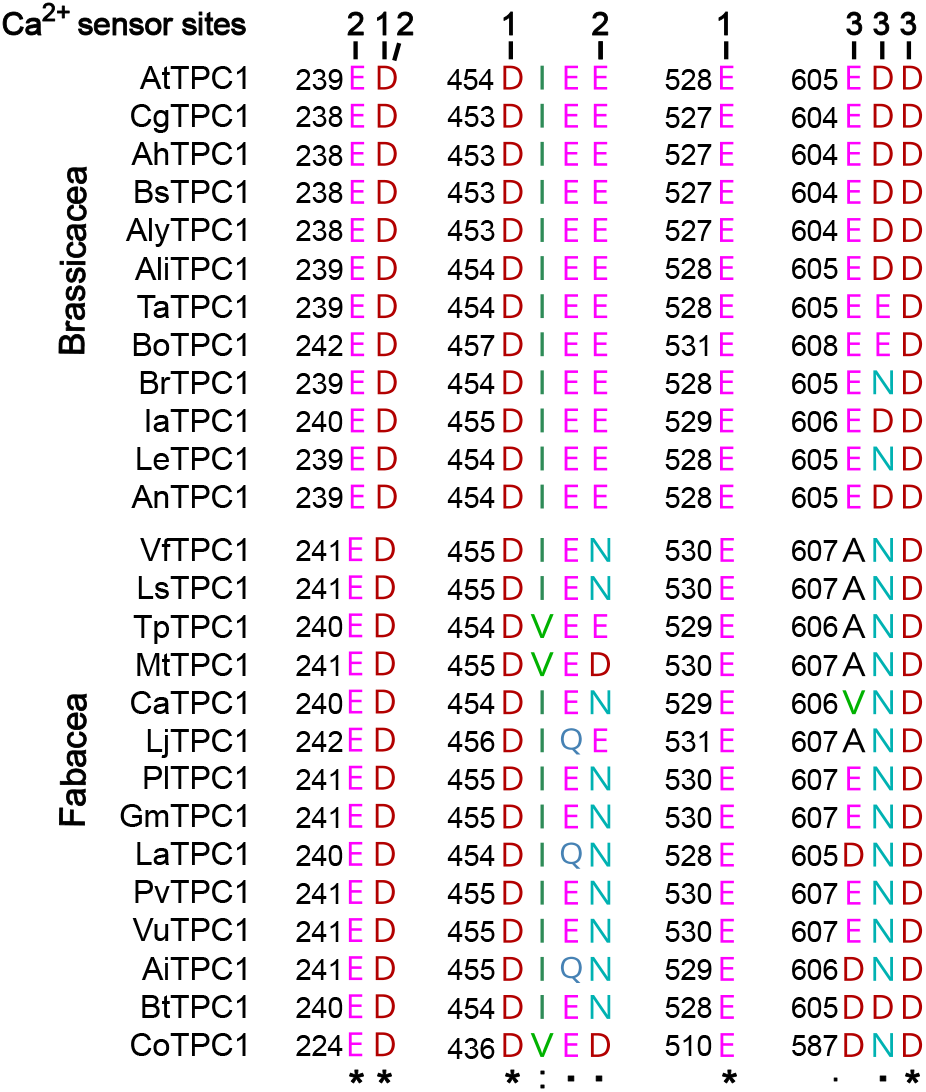
Polymorphism of functional TPC1 channel domains with a role in luminal Ca^2+^ coordination. Sections of an amino acid sequence alignment of Brassicacea and Fabacea TPC1 channels in the region of the three Ca^2+^ sensor sites. The numbers above the residues of AtTPC1 indicate to which Ca^2+^ sensor site they contribute. A different color code was used for different amino acids. An asterisk marks 100% conserved residues across the sequences while a colon (:) indicates a conservative and a dot (.) denotes a non-conservative substitution. TPC1 sequences from following species were used: *Arabidopsis thaliana* (AtTPC1), *Capsella grandiflora* (CgTPC1), *Arabidopsis halleri* (AhTPC1), *Boechera stricta* (BsTPC1), *Arabidopsis lyrata* (AlyTPC1), *Alyssum linifolium* (AliTPC1), *Thlaspi arvense* (TaTPC1), *Brassica oleracea* (BoTPC1), *Brassica rapa* (BrTPC1), *Iberis amara* (IaTPC1), *Lepidium sativum* (LeTPC1), *Arabis nemorensis* (AnTPC1), *Vicia faba* (VfTPC1), *Lathyrus sativus* (LsTPC1), *Trifolium pratense* (TpTPC1), *Medicago truncatula* (MtTPC1), *Cicer arietinum* (CaTPC1), *Lotus japonicus* (LjTPC1), *Phaseolus lunatus* (PlTPC1), *Glycine max* (GmTPC1), *Lupinus albus* (LaTPC1), *Phaseolus vulgaris* (PvTPC1), *Vigna unguiculata* (VuTPC1), *Arachis ipaensis* (AiTPC1), *Bauhinia tomentosa* (BtTPC1), *Copaifera officianalis* (CoTPC1). Amino acid sequences and their sources are listed in File S2.

### Polymorphic AtTPC1 triple mutant mimics the VfTPC1 channel features

To clarify whether these three polymorphic sites are responsible for the different gating behavior of the two TPC1 channel variants, a series of site-directed mutagenesis was initiated in AtTPC1. This resulted in the double and triple AtTPC1 mutants E605A/D606N and E457N/E605A/D606N. When transiently expressed in the background of the *tpc1-2* mutant (Fig. S1), these channel mutants gave rise to the typical SV/TPC1 channel currents (Fig. 1A). In the absence of luminal Ca^2+^, the current voltage curves (*I*_ss_/*C*_m_(*V*)) and even more so the activation voltage curves (*G*/*G*_max_(*V*)) of the double and triple mutants appeared shifted to more negative voltages by about 30 mV compared to wild type AtTPC1 (Fig. 1b, 2A and S4B). The changed voltage dependence of both AtTPC1 mutants was reflected in *V*_1_ and *V*_2_ values similar to wild type VfTPC1 (Fig. 2B). When the two mutants faced 10 mM luminal Ca^2+^, again the current and activation voltage curves as well as the *V*_1_, *V*_2_ values resembled those of VfTPC1 (Figs. 1B and 2). The fact that AtTPC1 channels carrying the VfTPC1 polymorphisms (i.e. double or triple residue substitutions) exhibit a low luminal Ca^2+^ susceptibility is best displayed by the channel behavior to 50 mM Ca^2+^. Under such high luminal Ca^2+^ loads, almost no SV/TPC1 currents were observed with wild type AtTPC1 (Fig. 1). The current and activation voltage curves (*I*_ss_/*C*_m_(*V*), *G*/*G*_max_(*V*)) of the AtTPC1 mutants and VfTPC1, however, showed similar strong SV channel activity even at this extreme luminal Ca^2+^ level (Fig. 1B and 2). Thus, the E605A/D606N exchange is already sufficient to provide AtTPC1 with the voltage and Ca^2+^ sensitivity of VfTPC1. An additional E457N replacement did not further affect the transition from an Arabidopsis-like to a VfTPC1-like gating behavior. Considering the attenuating effect of neutralized site 457 (E → N/Q) on voltage-dependent activation of AtTPC1 (12), the hyperactivity of the native *Vicia faba* TPC1 channel is most likely related to the altered Ca^2+^ sensor site 3.

### 3D topology of VfTPC1 and AtTPC1 triple mutant mimics that of *fou2*

In an attempt to determine the structural basis for the functional differences between *V. faba* and *A. thaliana* TPC1, we aimed to determine the cryoEM structure of VfTPC1, expressed and purified by similar conditions to that of AtTPC1 (16). Surprisingly, the biochemical behavior of VfTPC1 was significantly different than that of AtTPC1 and we were unable to recover any usable material after purification. Hence, we constructed a homology model of VfTPC1 using our high resolution cryoEM structure of Ca^2+^-bound AtTPC1-D454N (*fou2*) as a template. From our previous structures of wild-type AtTPC1 and *fou2* (14, 16), we determined that the pore mouth operates a luminal, inhibitory Ca^2+^ sensor (site 3) that is coupled to the functional voltage sensing domain (VSD2). In the wild-type and *fou2* structures, the three luminal acidic residues (E605, D606 and D607) of site 3 line the conduction pathway upstream of the selectivity filter. In the wild-type crystal structure, E605 binds a divalent metal on the symmetry axis. In our high-resolution structure of *fou2* (16), we showed that these residues rearrange to interact with the hydration shell around a bound metal in the selectivity filter and that D606 in *fou2* moves to the position of E605 in the wild-type structure, suggesting that both acidic residues regulate channel function. In our homology model of VfTPC1, neutralization of pore mouth acidic residues A607 and N608 (equivalent to E605 and D606 in AtTPC1) clearly alters the electrostatics of the pore. These substitutions almost certainly prevent the pore mouth from binding the inhibitory Ca^2+^ observed in the wild-type structure (Fig. 4A), and from interacting with the water network in the pore as seen in *fou2* (16). Recent molecular dynamics (MD) simulations with AtTPC1 (25) additionally indicated that the E605/D606 motif is structurally very flexible and adapts according to the direction of ion flow. This may suggest a valve-like regulatory mechanism: The efflux from the cytosol to the lumen stabilizes the position found in the hyperactive *fou2* mutant, *i.e.*, the open channel. In contrast, influx from the lumen to the cytosol drives the E605/D606 loop towards the pore, fostering channel closing. The MD simulations further suggest a Ca^2+^-dependent interaction between the E605/D606 motif and a Ca^2+^ coordination site close to the luminal entrance of the selectivity filter (D269/E637; in VfTPC1: D271/E639). Higher luminal Ca^2+^ increases the probability of Ca^2+^ to occupy the coordination site which stabilized the E605/D606-motif in the pore-facing conformation. Neutralizing E605A/D606N mutations would reduce electrostatic interactions and result in channels that open with less activation energy.

**Fig. 4.**
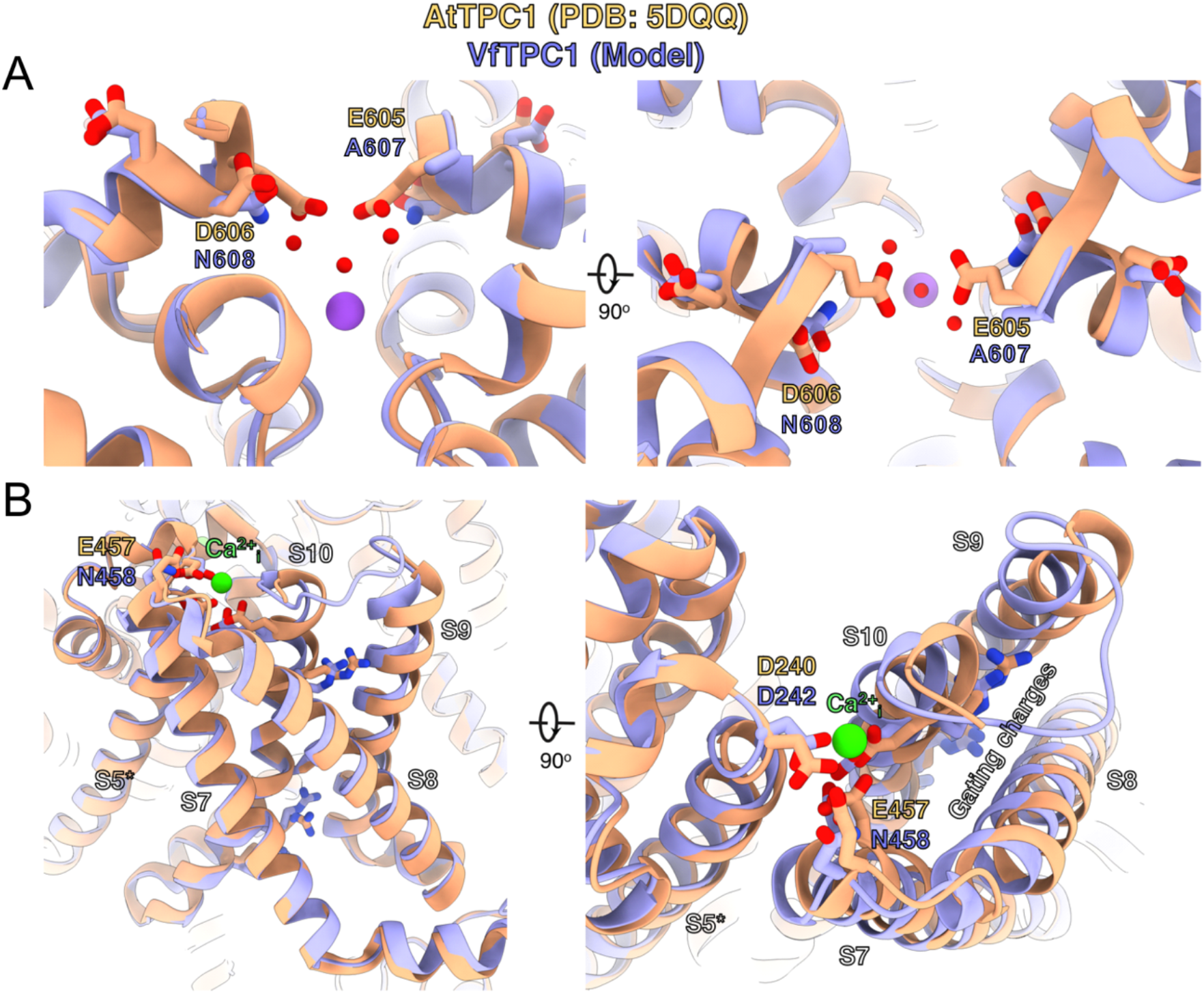
Structural comparison between wild type AtTPC1 and a VfTPC1 homology model depicts functional consequences of sequence divergence. (*A*) Orthogonal views of the pore mouth residues. The central purple and red spheres are Na^+^ and waters, respectively, as observed in the cryoEM structure of *fou2* (16). (*B*) Orthogonal views of the luminal Ca^2+^-binding site on VSD2. For all panels, AtTPC1 is beige and VfTPC1 purple.

### Mutation of AtTPC1 pore motif E605/D606 does neither affect SV channel permeability nor conductance

The pore residues E605 and D606 are located relatively close to the selectivity filter (Fig. 4; 26). Therefore, we investigated their contribution to the TPC1 cation permeability (Na^+^, K^+^, Ca^2+^) in the animal HEK cells that target TPC1 to the plasma membrane (Fig. S6; 13). To enable current recordings with AtTPC1 even under high external Ca^2+^ conditions, the AtTPC1 channel mutant D240A/D454A/E528A (27) harboring a damaged luminal Ca^2+^ sensing site 1 in the presence of a wild-type pore region was used. The relative permeability ratio P_K/Ca_/P_Na_ of AtTPC1 was not affected by disruption of site 1 (D240A/D454A/E528A; 27) or site 3 (E605A/D606A) (Table S2). In comparison to the AtTPC1 channel variants, VfTPC1 was characterized by a similar cation permeability. Thus, structural domains other than the pore motif E605/D606 determine the TPC1 channel selectivity.

BK channels are Ca^2+^-activated K^+^ channels with large conductance, similar to TPC1 (28). A negative charge ring at the pore entrance promotes the single channel conductance of BK channels (29). To examine whether the negatively charged residues at the luminal pore entrance of AtTPC1 may play a similar role as in BK channels, single channel currents were recorded (Fig. S7A). Under symmetrical K^+^ conditions (100 mM) a single channel conductance of about 80 pS was determined for AtTPC1 wild type (Fig. S7B and C). With respect to the dimeric structure of TPC1, the neutralizations in the AtTPC1 double mutant E605A/D606A results in the removal of a total of four negative charges in the luminal pore vestibule. Nevertheless, the unitary conductance of the mutant remained unchanged. Moreover, VfTPC1 has a threefold higher unitary conductance than AtTPC1 (250 pS, Fig. S7; cf. 30), despite its neutral luminal pore residues at the homologous sites to AtTPC1 (E605, D606). These facts indicate that these luminal polymorphic pore residues of Ca^2+^ sensor site 3 apparently do not contribute to the conductance of TPC1.

### VfTPC1 and AtTPC1 triple mutant are hyperexcitable

The increased activity of the AtTPC1 channel mutant *fou2* leads to a hyperexcitability of the vacuole (5). Here, the membrane polarization was measured at physiological 0.2 mM luminal Ca^2+^ (31) with vacuoles equipped with either VfTPC1, AtTPC1 wild type or the triple AtTPC1 mutant E457N/E605A/D606N. For TPC1-dependent vacuolar excitation, depolarizing current pulses of increasing amplitudes in the range of 10 to 1000 pA were temporarily injected from a resting voltage of −60 mV (Fig. 5; S8A-C; cf. 5). After the depolarizing current stimulus (Fig. 5A and S8A-C), the vacuole membrane equipped with VfTPC1 channels remained depolarized at a voltage of about 0 mV (i.e. at the equilibrium potential for K^+^) during the entire subsequent recording period of 10 s, regardless of the stimulus intensity (Fig. 5, Table S1). In contrast, when AtTPC1-equipped vacuoles were challenged with even the highest current stimulus (1 nA), the post-stimulus voltage remained depolarized for only a short period (*t*_plateau_ ~ 0.4 s, Table S1) before relaxation to the resting voltage occurred. In comparison to AtTPC1 wild type, the presence of the AtTPC1 triple mutant in the vacuole membrane strongly prolonged the lifetime of the depolarized post-stimulus voltage-plateau phase at all current stimuli (Fig. 5, Table S1). Except of one vacuole, all other five AtTPC1-triple mutant vacuoles already responded to the lowest current pulses such as 10, 30 or 70 pA with sustained post-stimulus depolarization over the entire recording period (Table S1). In other words, AtTPC1-triple mutant vacuoles were also hyperexcitable, behaving very much like the VfTPC1 vacuoles. This points to a correlation between the vacuolar hyperexcitability and the shifted voltage activation threshold of the hyperactive TPC1 channels (Figs. 5, S8).

**Fig. 5.**
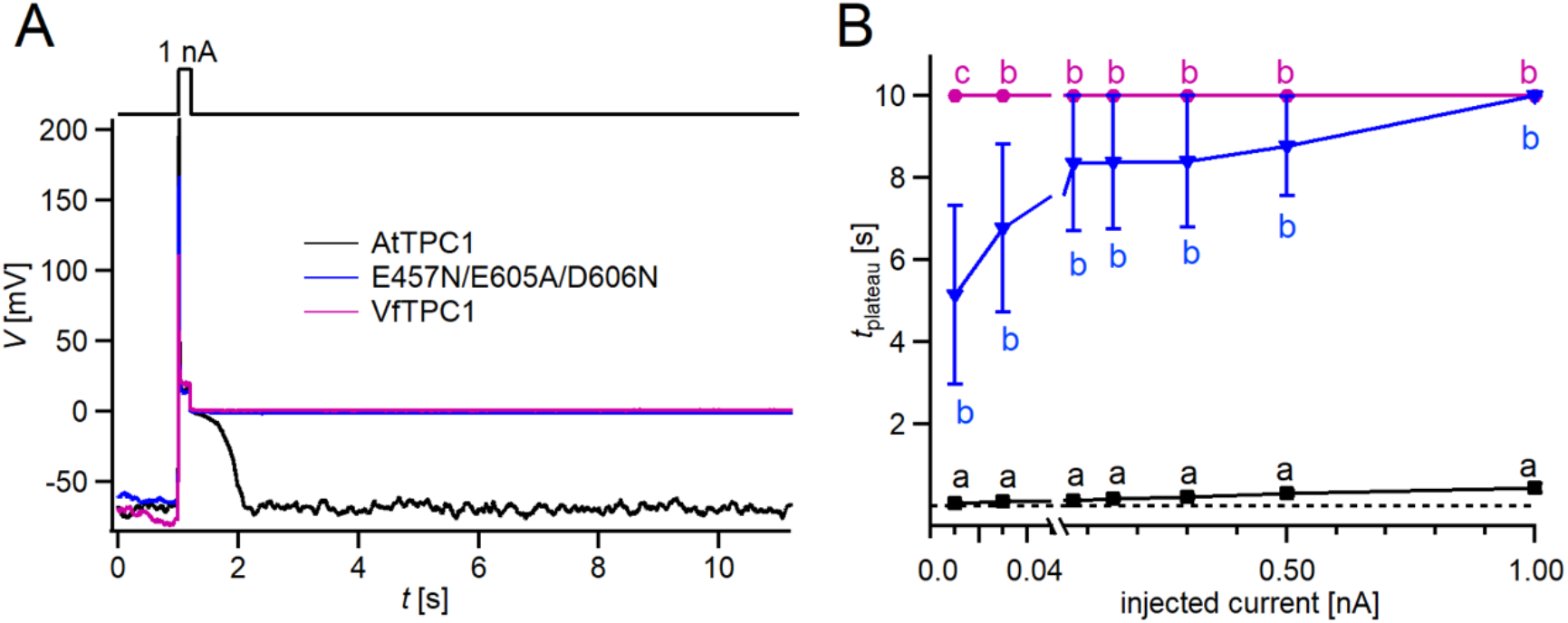
Dependency of vacuole excitability on TPC1 channel variants. (*A*) Superimposed voltage responses (lower panel) of individual vacuoles equipped either with VfTPC1 wild type (magenta), AtTPC1 wild type (black) or AtTPC1-triple mutant E457N/E605A/D606N (blue) to current injection of 1 nA (upper panel). (*B*) Lifetime of the post-stimulus depolarization phase plotted against the corresponding injected current pulse. Symbols represent means ± SE (squares = AtTPC1 wild type; circles = VfTPC1 wild type; reversed triangles = AtTPC1-E457/E605A/D606N). Individual data points are listed in Table S1. Number of experiments for each channel type was n = 6. Significant differences between the channel variants are indicated by different letters (one-way ANOVA followed by a Tukey’s post hoc comparison test). All experiments were carried out at 0.2 mM luminal Ca^2+^.

In the following we used a computational model to simulate activation of voltage-dependent TPC1-mediated vacuole excitability (5) (Fig. 3). Under standard conditions (Fig. 6, blue) a current stimulus induced a typical excitation as observed in whole-vacuole patch clamp experiments (Figs. 5 and 6B). In the simulation too, the duration of the post-stimulus voltage-plateau phase (*t*_plateau_) was found to depend on the stimulus strength (Fig. 6C and E). The voltage activation threshold of TPC1 had an additional influence on *t*_plateau_. A negatively shifted TPC1 activation curve (Fig. 6A, black) provoked that the vacuole membrane remained in the excited state after stimulation (Fig. 6B, black). A positively shifted TPC1 activation curve (Fig. 6A, pink), however, shortened the plateau phase and thus of the time course of the excited state (Fig. 6B and D, pink). As a consequence, the slope of the linear relationship between *t*_*plateau*_ and stimulus strength was reduced (Fig. 6E). Thus, effectors regulating the voltage activation threshold of TPC1 (e.g., Ca^2+^) tune the sensitivity of the vacuole membrane and shape the duration of the excited state. In comparison to AtTPC1 expressing vacuoles, those equipped with VfTPC1 or the AtTPC1 triple mutant, which both activate more negatively, are hyperexcitable (Fig. 5).

**Fig. 6.**
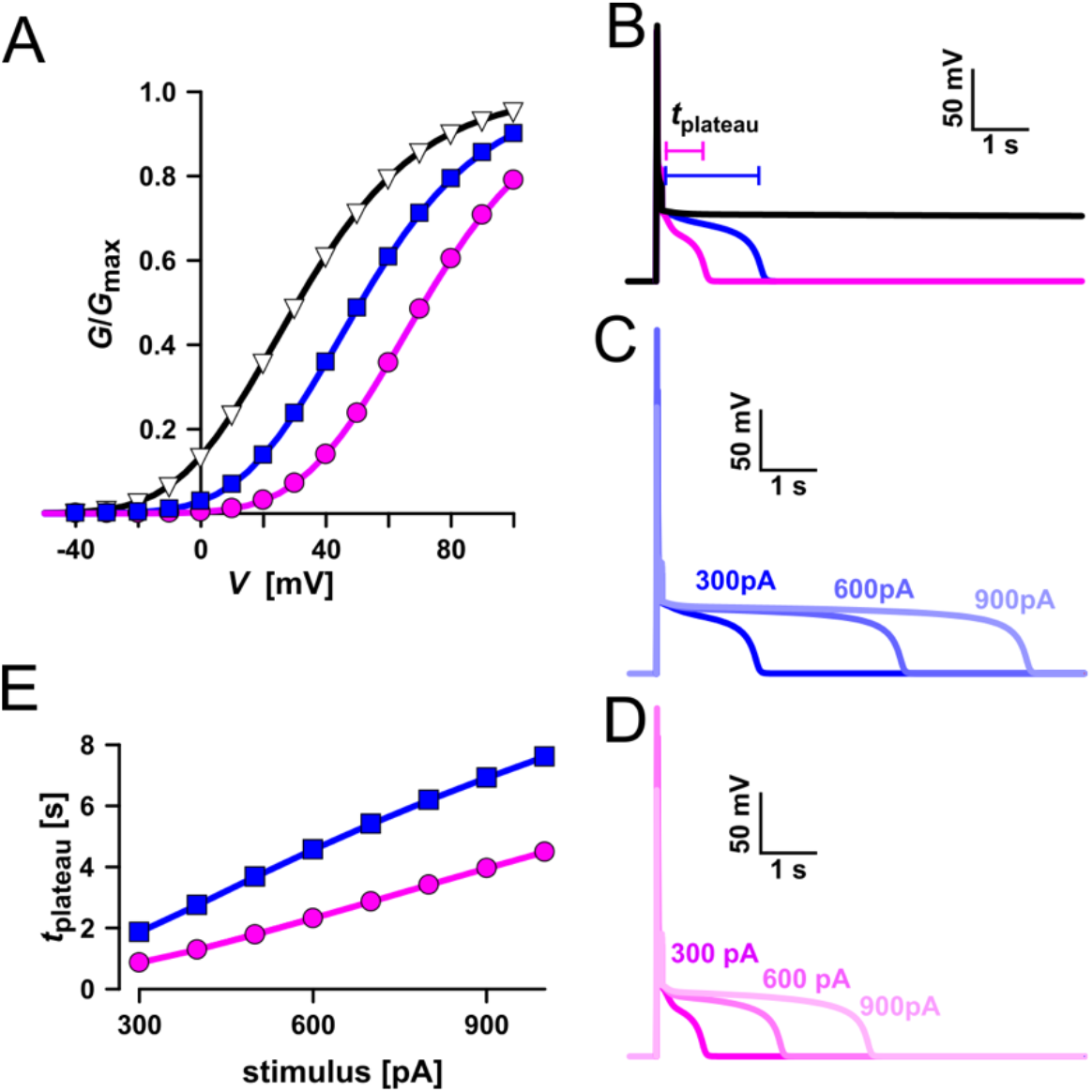
Simulation of vacuolar electrical excitability with three different TPC1 variants. (*A*) Gating characteristics (*G*/*G*_max_) of three different TPC1-type channels. (*B*) Simulation of vacuolar electrical excitability with the different TPC1 variants. Excitation was induced by a 300 pA pulse of 100 ms duration. (*C*) Overlay of the electrical response of the vacuole having the blue-type TPC1 to 100 ms pulses of 300 pA, 600 pA and 900 pA. (*D*) Overlay of the electrical response of the vacuole having the red-type TPC1 to 100 ms pulses of 300 pA, 600 pA and 900 pA. (*E*) Dependency of the length of the post-stimulus plateau phase on the stimulus strength.

## Discussion

We identified the Fabaceae VfTPC1 channel from *Vicia faba* as the first natural hyperactive TPC1 channel variant. The VfTPC1 hyperactivity results from the activation near vacuolar resting membrane voltage and the luminal Ca^2+^ insensitivity (Fig. 2). The dual polymorphism of two residues within the Ca^2+^ sensor site 3 in the luminal pore entrance is most likely responsible for this gating behavior of VfTPC1, which is clearly distinct from the AtTPC1 channel of Brassicaceae. In (16) we proposed that Ca^2+^ sensor site 3 in AtTPC1 is formed by three negatively charged residues (E605, D606, D607). Here, we have now found that E605 and D606 play the most important role within this triad of site 3 in gating control (Fig. 2). Neutral residues at this site, as naturally already present in VfTPC1 (A607/N608; Fig. 3), uncouple luminal Ca^2+^ coordination from gating, leading to channel hyperactivity (Fig. 2).

### VfTPC1 versus AtTPC1: what is the rule and what is the exception?

*Arabidopsis thaliana* is the model plant for plant molecular genetics and physiology. Is the AtTPC1 structure-function-relation reference for all plant SV channels? Obviously not! Unlike AtTPC1, VfTPC1 is hyperactive and less sensitive towards luminal Ca^2+^ (Figs. 1, 2, 4 and S4). In fact, these VfTPC1 features may be even more widespread in the plant kingdom, because TPC1 channels from some other plant species like e.g. *Lotus japonicus, Oryza sativa* and *Hordeum vulgare* also exhibit the neutralizing triple-polymorphism characteristic of VfTPC1 in the luminal Ca^2+^ sensor sites 2 and 3 (Figs. 3, S9). In well agreement with this triple polymorphism, the fabacean TPC1 channel from *Lotus japonicus* shows a VfTPC1-like gating behavior (Fig. S10). Similarly, voltage- and Ca^2+^-dependent TPC1 current responses from mesophyll vacuoles of the monocots *O. sativa* and *H. vulgare* appear also to exhibit a lower luminal Ca^2+^ sensitivity and higher probability to open at more negative voltages, respectively (21, 23, 32). Such luminal pore polymorphism, feeding back on gating behavior of TPC1 channels, could allow species to adapt the electrical properties of the vacuole to given individual settings.

### The complex relationship between TPC1 and Ca^2+^

In well agreement with experiments with *Vicia faba* vacuoles (22, 33), our cation replacement experiments with TPC1-expressing HEK cells (Fig. S6, Table S2) demonstrate the principal ability of VfTPC1 and AtTPC1 to conduct not only K^+^ and Na^+^ but also Ca^2+^ ions to similar ratios under these unphysiological settings. However, under physiological, plant cell-like Ca^2+^ and K^+^ concentrations and gradients, the fava bean SV/TPC1 channel was predominately conducting K^+^ (30, for review see 2, 34). The MD simulations with AtTPC1 also demonstrated K^+^ and Na^+^ but not Ca^2+^ permeation (25). This permeation profile is consistent with the K^+^-starvation transcriptome and the elevated vacuolar Ca^2+^ content of Arabidopsis plants equipped with hyperactive *fou2* TPC1 channels (20, 35). Thus, these observations contradict the idea that TPC1 may operate as <u>C</u>a^2+^-induced <u>C</u>a^2+^ release (CICR) element in the vacuole membrane (22). Instead, TPC1-triggered vacuole excitation can cause the release of Ca^2+^ from the vacuole to the cytosol via a Ca^2+^ homeostat with a striking correlation between the duration of excitation and the amount of released Ca^2+^ (36). Given that the voltage activation threshold of TPC1 is under control of luminal Ca^2+^ (Fig. 2), it directly influences the excitation duration of the vacuole membrane (Figs. 4, 5 and 6E) and Ca^2+^ release in turn. Without such a regulatory process, a higher luminal Ca^2+^ concentration would imply a larger transmembrane Ca^2+^ gradient and therefore release a larger Ca^2+^ quantum. Tuning of the TPC1 activity by luminal Ca^2+^ has the potential to mitigate or even counterbalance the electro-chemical potential of a larger Ca^2+^ gradient. A higher luminal Ca^2+^ concentration downregulates the TPC1 activity, which in turn reduces the excitation duration and results in a shortened Ca^2+^ release of larger amplitude. The feedback mechanism on gating thus largely uncouples the Ca^2+^ release mechanism from the luminal Ca^2+^ content and keeps it operational under a broad range of environmental-induced Ca^2+^ signaling scenarios.

In view of the high impact of the TPC1 activation threshold on hormone homeostasis and plant growth, future research is now warranted to answer the question why *Vicia faba* plants do not exhibit a retarded growth phenotype, as Arabidopsis mutant plants with the artificial hyperactive AtTPC1 channel *fou2* do (19).

## MATERIALS AND METHODS

### Plant materials and growth conditions

The *Arabidopsis thaliana tpc1-2* mutant (4) was grown in a growth chamber under short day conditions (8 h light, 16 h dark) with a day/night temperature regime of 22/16 °C, a photon flux density of 150 μmol m^−2^ s^−1^ and a relative humidity of about 60%. *Vicia faba* and *Lotus japonicus* plants were grown in the greenhouse at a 16 h day/8 h dark photoperiod, a day/night temperature regime of 22/18°C and a light intensity of approximately 1250 μmol m^−2^ s^−1^.

### RNA sequencing

Leaves from 6- to 8-week-old *Vicia faba* plants were harvested and mesophyll RNA was extracted using the NucleoSpin Plant RNA extraction kit (Macherey-Nagel, Düren, Germany) according to the manufacturer’s instructions. For guard cell RNA extraction epidermal fragments were isolated using the blender method (37, 38). Total RNA from three individual biological replicates was prepared and subjected to RNA-sequencing on an Illumina NextSeq500 platform. High-quality RNA-seq paired-end reads were quality checked using FastQC (version 0.11.6) and transcriptomes were *de novo* assembled individually using Trinity (version 2.5.1 Release). Finally, the TRAPID pipeline was employed for processing of assembled transcriptome data including transcript annotation (39). Based on AtTPC1 homology, identical VfTPC1 transcripts were identified in mesophyll and guard cell fractions, and the obtained sequence information was used to clone the VfTPC1 CDS by a PCR-based approach. The *VfTPC1* mRNA sequences is deposited at Genbank under the following accession number, respectively: BankIt2410619 VfTPC1_mRNA MW380418.

### Cloning and site-directed mutagenesis

After the total RNA was extracted from mature leaves of *Vicia faba* and *Lotus japonicus* plants and reverse transcribed into complementary DNA (cDNA), VfTPC1 and LjTPC1 were amplified from the cDNA libraries, essentially as described (23). For patch clamp experiments with vacuoles, the cDNA coding sequences of the AtTPC1 channel variants were cloned into the modified pSAT6-eGFP-C1 vector (GenBank AY818377.1), whereas the coding sequences of the VfTPC1 and LjTPC1 channel variants were cloned into the pSAT6-eYFP-C1 vector (GenBank DQ005469) using the uracil-excision-based cloning technique (40), essentially as described by (12). The resulting AtTPC1-eGFP constructs were under the control of the 35S promoter, and the VfTPC1-eYFP and LjTPC1-eYFP constructs were under the control of the ubiquitin promoter (UBQ10). For patch clamp experiments with HEK cells, the eYFP coding sequence (41) was fused without the stop codon to the 5’ end of the TPC1 cDNA coding sequences and then cloned together into the pcDNA3.1 vector (GenBank MN996867.1). In analogy to (12), a modified USER fusion method was used to introduce site-directed mutations in the wild type AtTPC1 construct. The sequences of the primers used for subcloning and mutagenesis are listed in Table S2. All channel variants were tested for their sequences.

### Transient protoplast transformation

Essentially following the protocols from (42) and (43), mesophyll protoplasts were released from 6- to 7-week-old *tpc1-2* Arabidopsis mutant plants and transiently transformed with the different TPC1 channel constructs. For channel expression, protoplasts were then stored in W5 solution (125 mM CaCl_2_, 154 mM NaCl, 5 mM KCl, 5 mM glucose, 2 mM Mes/Tris, pH 5.6, 50 μg ml^−1^ ampicillin) at 23°C in the dark for usually two days.

### HEK cell transfection

HEK293 cells were transfected with the respective eYFP-TPC1-fused constructs with lipofectamine2000 (ThermoFisher, Waltham, USA) according to the manufacturer’s instructions. Cells were seeded 18-24 hours after transfection in glass coverslips (diameter 12 mm). Protein expression was verified on the single-cell level by the appearance of eYFP-fluorescence in the plasma membrane upon excitation with a 473 nm laser.

### Subcellular targeting

Vacuolar membrane localization of the expressed eGFP/eYFP-fused TPC1 channels was verfied by imaging the fluorescence signal of transformed protoplasts and vacuoles with a confocal laser scanning microscope (TCS SP5, Leica, Mannheim, Germany) (23). eGFP and eYFP were excited with an Argon laser at 490 and 514 nm, respectively, and the emission of fluorescence was monitored between 500 and 520 nm for eGFP and between 520–540 nm for eYFP. Red autofluorescence of chlorophyll was excited at 540 nm and acquired between 590 and 610 nm. For expression analysis in HEK293 cells, a confocal laser scanning microscope (SP700, Zeiss, Germany) equipped with three laser lines (488 nm: 10 mW, 555 nm: 10 mW, 639 nm: 5 mW) was used. Images were processed with ZEN software (ZEN 2012, Zeiss) or Fiji, Version ImageJ 1.50 (44).

### Whole-vacuole patch clamp experiments

Vacuoles were released from protoplasts in the recording chamber two days after transformation using a vacuole release (VR) solution (45). The VR solution was modified and composed of 100 mM malic acid, 155 mM N-methyl-D-glucamine, 5 mM EGTA, 3 mM MgCl_2_, 10 mM Hepes/Tris pH 7.5 and adjusted to 450 mOsmol.kg^−1^ with D-sorbitol. The whole vacuole configuration was then established with TPC1-transformed vacuoles which were easily identified upon their eGFP- or eYFP-based fluorescence measured between 510 and 535 nm after excitation at 480 nm with a *precisExcite HighPower* LED lamp (Visitron Systems GmbH, Puchheim, Germany). Patch pipettes were prepared from Harvard glass capillaries (Harvard glass capillaries GC150T-10, Harvard Apparatus, UK) and typically had a resistance in the range of 1.4-3.1 MΩ. The membrane capacitance (*C*_m_) of the individual vacuoles accessed and compensated in patch clamp experiments ranged from 31.1 to 68.5 pF. Membrane currents and voltages were recorded with a sampling rate of 150 μs at a low pass filter frequency of 2.9 or 3 kHz using an EPC10 or EPC800 patch clamp amplifier, respectively (HEKA Electronic). Data were acquired with the software programs Pulse or Patchmaster (HEKA Electronic) and off-line analyzed with IGORPro (Wave Metrics). Voltage recordings were carried out in the current-clamp mode as described (5). Briefly, after adjusting the membrane voltage to −60 mV by injection of an appropriate current, current pulses were applied in the range of 10 to 1000 pA for 200 ms. The duration of the post-stimulus depolarisation phase gives the time at which the initial depolarized voltage dropped by 50% to the holding voltage.

In voltage-clamp experiments macroscopic currents were recorded in response to 1-s-lasting voltage pulses in the range of −80 mV to +110 mV in 10 mV increments. The holding voltage was −60 mV. The corresponding current responses of each vacuole were analyzed with respect to the half-activation time (*t*_act-0.5_) and steady-state current amplitudes (*I*_ss_). The half-activation time (*t*_act-0.5_) was the time at which 50% of the steady-state current amplitude was reached. As a normalization measure for the membrane surface of the individual vacuole, the determined steady state currents were divided by the respective compensated membrane capacitance (*C*_m_). Conductance/voltage curves (*G/G*_max_(*V*)) were quantified from tail current experiments as a measure for the relative voltage-dependent open channel probability. Following pre-pulse voltages in the range of −80 to +110 mV, instantaneous tail currents were determined at −60 mV. The midpoint voltages (*V*_1_, *V*_2_) and the equivalent gating charges (*z*_1_, *z*_2_) were derived by fitting the *G/G*_max_(*V*) curves with a double Boltzmann equation (16). After pre-activation of TPC1 currents upon an instantaneous voltage pulse to either +80 mV or +100 mV, the current relaxation was recorded at voltages from −60 to 0 mV and the deactivation-half times (*t*_deact-0.5_) were determined. The half-deactivation time (*t*_deact-0.5_) denotes the time at which the initial tail current amplitude has declined by 50%.

In voltage-clamp experiments the standard bath solution facing the cytoplasmic side of the vacuole membrane contained 150 mM KCl, 1 mM CaCl_2_, 10 mM Hepes (pH 7.5/Tris) and was adjusted with D-sorbitol to an osmolality of 520 mOsmol.kg^−1^. The standard pipette solution at the luminal side of the tonoplast basically consisted of 150 mM KCl, 2 mM MgCl_2_, 10 mM HEPES (pH 7.5/Tris) and was adjusted with D-sorbitol to a final osmolality 500 mOsmol.kg^−1^. The pipette solution was supplemented with 10 mM or 50 mM CaCl_2_ or adjusted to 0 mM Ca_2_ by addition of 0.1 mM EGTA. In current-clamp experiments the standard bath medium additionally contained 2 mM MgCl_2_ and the pipette solution was adjusted to 0.2 mM free Ca^2+^ by addition of 4.1 mM EGTA and 4.3 mM Ca^2+^ (https://somapp.ucdmc.ucdavis.edu/pharmacology/bers/maxchelator/webmaxc/webmaxcE.htm).

### Patch clamp experiments with membrane patches

Vacuoles were isolated from transiently transformed protoplast by perfusion with a solution containing 10 mM EGTA, 10 mM Hepes/Tris pH 7,5, adjusted to 200 mosmol kg^−1^ with D-Sorbitol. Excised membrane patches with the cytosolic side of the tonoplast facing the bath medium were formed from the whole vacuole configuration. Single channel fluctuations were recorded with a sampling rate of 100 μs at a low pass filter frequency of 1 kHz using an EPC10 patch clamp amplifier. Single channel current amplitudes were determined from all-point histograms of the current recordings and plotted against the respective voltages. The single channel conductance for each membrane patch was derived from linear regression of the current-voltage plot. The bath medium contained 100 mM KCl, 0.5 mM CaCl_2_, 10 mM Hepes (pH 7.5/Tris). The pipette medium consisted of 100 mM KCl, 2 mM MgCl_2_, 2 mM EGTA, 10 mM MES (pH 5.5/Tris). Both solutions were adjusted to 400 mosmol.kg^−1^ with sorbitol.

### Patch clamp experiments with HEK cells

Whole-cell current recordings were performed at a setup described previously (46). Data were acquired using Clampex 10.7 (Molecular devices, San Jose, USA) with 100 kHz sampling rate, low-pass filtered at 5 kHz, and analyzed with Clampfit 10.7 (Molecular devices) and OriginPro 2016 (Originlab, Northampton, USA). Pipette resistance (GB150F-8P, Scientific-Instruments, Hofheim, Germany) was 4-6 MΩ in standard bath solution. The standard pipette solution contained 150 mM NaCl, 2.5 mM MgCl_2_, 0.3 mM free Ca^2+^ (adjusted by 4.3 mM CaCl_2_ and 4 mM EGTA), 10 mM Hepes, (pH 7.4/Tris). The standard bath solution was composed of 150 mM NaCl, 10 mM HEPES, (pH 7.4/Tris). Under these experimental conditions bath and pipette solutions mimicked the vacuolar and cytosolic solute conditions, respectively. To determine the relative permeability ratio on the same HEK cell under bi-ionic cation conditions, the Na^+^-based bath medium was replaced by either a K^+^- or Ca^2+^-based solution. The K^+^-based bath medium contained 150 mM KCl, 10 mM HEPES, (pH 7.4/Tris), and the Ca^2+^-based one consisted of 15 mM CaCl_2_, 120 mM NMDG-Cl, 10 mM Hepes (pH 7.4/Tris).

For determination of the TPC1 ion selectivity, tail current experiments were conducted. After pre-activating the TPC1 channels by a voltage step from −70 mV (resting potential) to +100 mV for 500 to 1000 ms, the relaxation of the outward currents in response to hyperpolarizing voltage pulses was recorded either with 10 mV intervals reaching from +60 mV to −120 mV or in 5 mV decrements reaching from +30 mV to −40 mV to visualize and to analyze the reversal potentials, respectively. In order to clearly assign the reversal potential to TPC1 currents, which are not contaminated by leakage currents, the slope of the tail currents was determined and plotted against the corresponding voltages. The reversal potential for each solute condition was then determined by interpolation the smallest negative and positive slope values. The shift in the reversal potential ΔV_rev_ caused by the change of the external solution from either Na^+^ to K^+^ or Na^+^ to Ca^2+^ solution was used to estimate the relative permeability ratios P_Ca_/P_Na_ or P_K_/P_Na_ essentially as described by (47). For calculation of the K^+^ to Na^+^ permeability ratio (P_K_/P_Na_) the following equation (1) was used (48):

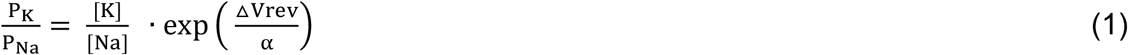

where ΔV_rev_ = V_K_-V_Na_, with the reversal potentials V_K_ and V_Na_ in K^+^- or Na^+^-based bath solution, respectively, and α= RT/F = 25.42 mV at 22°C. The relative permeability ratio P_Ca_/P_Na_ was calculated using equation (2) derived from the extended Goldman-Hodgkin-Katz equation (49):

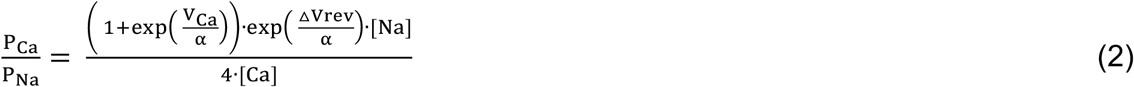

where ΔV_rev_ = V_Ca_-V_Na_ with the reversal potentials V_Ca_ and V_Na_ in Ca^2+^ - or Na^+^-based bath solution, respectively, and α has same meaning as above.

### Voltage convention in patch clamp experiments

The given membrane voltages refer to the cytosolic side of the vacuole membrane or HEK cell plasma membrane with zero potential on its luminal or extracellular side, respectively. In experiments with vacuoles performed in the presence of 50 mM luminal Ca^2+^ (Figs. 1, 2, S2D and S3D), the membrane voltages were corrected off-line by the corresponding liquid junction potential determined offline according to (50) and (22). Otherwise, no correction for the liquid junction potential was necessary.

### Statistical analysis

Patch clamp experiments were conducted with individual vacuoles per each channel construct and solute condition. Due to a common data set, *I*_ss_/*C*_m_(V) and *G/G*_max_(*V*) curves for wild type AtTPC1 acquired under 0 and 10 mM luminal Ca^2+^ were identical to those shown in (16). Electrophysiological data are given as means ± SE or SD as indicated in the figure legends. The statistical analysis was performed using one-way-ANOVA followed by the Dunnett’s or Tukey’s posthoc comparison test. In order to test also the Ca^2+^-induced shifts in the *V*_1/2_ values for significant differences, the *V*_1_ and *V*_2_ mean values determined in the absence of luminal Ca^2+^ were subtracted from the individual *V*_1_ and *V*_2_values, respectively, derived under 10 mM luminal Ca^2+^ conditions (16). In analogy, the Ca^2+^-induced change in the *t*_act-0.5_ values were analyzed for significant differences. Statistical analysis was done with Origin 2020 (OriginLab, Northampton, Massachusetts, USA) and SPSS 2020 (IBM, New York, USA).

### 3D modelling

An atomic model of VfTPC1 was generated using the homology modelling program MODELLER B. (51), using the experimental AtTPC1 D454N (*fou2*) Ca^2+^-bound structure (16) as a reference. The model was then relaxed into the 2.5 Å resolution *fou2*-Ca^2+^ map (16) using ISOLDE (52) and used for atomic interpretation.

### In silico experiments

Electrical excitability of the vacuolar membrane was computationally simulated as described in detail before (5) involving a background conductance, voltage-independent K^+^ channels of the TPK-type and the time- and voltage-dependent cation channel TPC1 that confers excitability to the vacuolar membrane. The delayed-activating behavior of the latter can be described mechanistically by four independent gates of two different types following the gating schemes *O*_1_ ↔ *C*_1_ and *O*_2_ ↔ *C*_2_ with the rate constants *a*_1_, *a*_2_, *d*_1_, and *d*_2_ for activation and deactivation, respectively: *a*_1_ = s^−1^ × exp[0.45 × *V* × *F*/(*RT*) - 0.26 × ln(α)], *d*_1_ = s^−1^ × exp[-0.81 × *V* × *F*/(*RT*) + 0.26 × ln(α) + 1.84], *a*_2_ = s^−1^ × exp[0.5 × *V* × *F*/(*RT*) – 0.26 × ln(α) – 0.4], *d*_2_ = s^−1^ × exp[-0.5 × *V* × *F*/(*RT*) + 0.26 × ln(α) + 3.0]. To simulate TPC1s with different gating features, the parameter has been set to 0.002, 0.01, and 0.05.

## Acknowledgements

The work was supported in part by a Reinhart Koselleck project (HE 1640/42-1; project number 415282803) from the German Research Foundation (DFG) to R.H., grants from the German Research Foundation (DFG) for the priority programs ‘MAdLand – Molecular Adaptation to Land: Plant Evolution to Change’ to R.H. and D.B, a doctoral fellowship from the China Scholarship Council (CSC) and a STIPET fellowship from the German Academic Exchange Service (DAAD) to J.L., the CONICYT-FONDEQUIP project EQM160063 to I.D. and C.N.-R., a Fondecyt-Enlace project of the Universidad de Talca to I.D., and the postdoc grant FONDECYT no. 3170434 of the Comisión Nacional Científica y Tecnológica of Chile to C.N.-R. We are grateful to Parathy Yogendran for transient transfection of HEK293 cells, Matthias Freund for his support in statistical analysis and Armando Carpaneto for discussion.

