## Supporting Information File S1 for "*Vicia faba* TPC1, a genetically encoded variant of the vacuole Two Pore Channel 1, is hyperexcitable"

### Supporting Information – File 1

### Figures, Tables and References

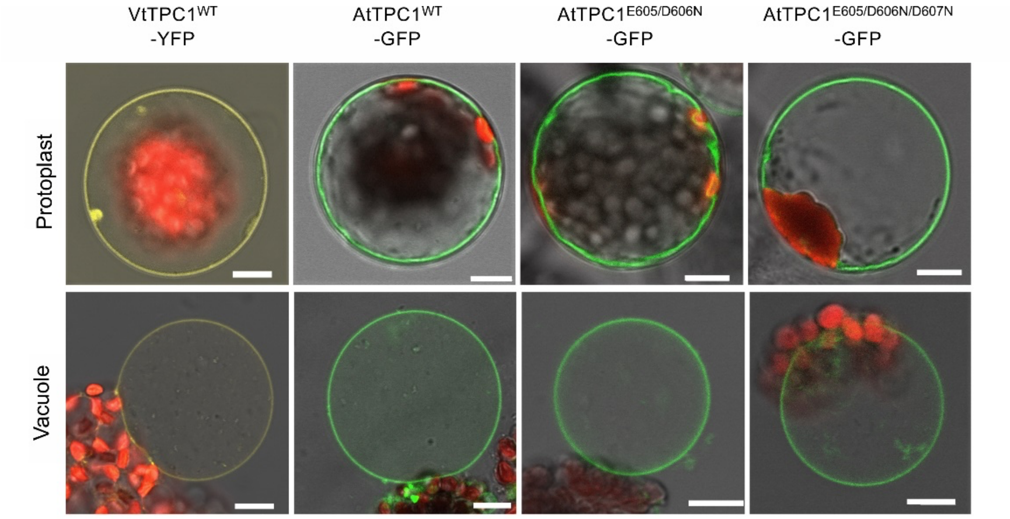


**Fig. S1.** Vacuolar targeting of TPC1 channel variants. Mesophyll protoplasts from the *Arabidopsis thaliana* mutant *tpc1-2* after transient transformation with the respective GFP/YFP-tagged TPC1 channel construct and released vacuoles. Bright field and fluorescent images were merged. Red fluorescence corresponds to chloroplast autofluorescence. The yellow and green fluorescence represent the YFP and GFP fluorescence, respectively. Scale bars = 10 μm.

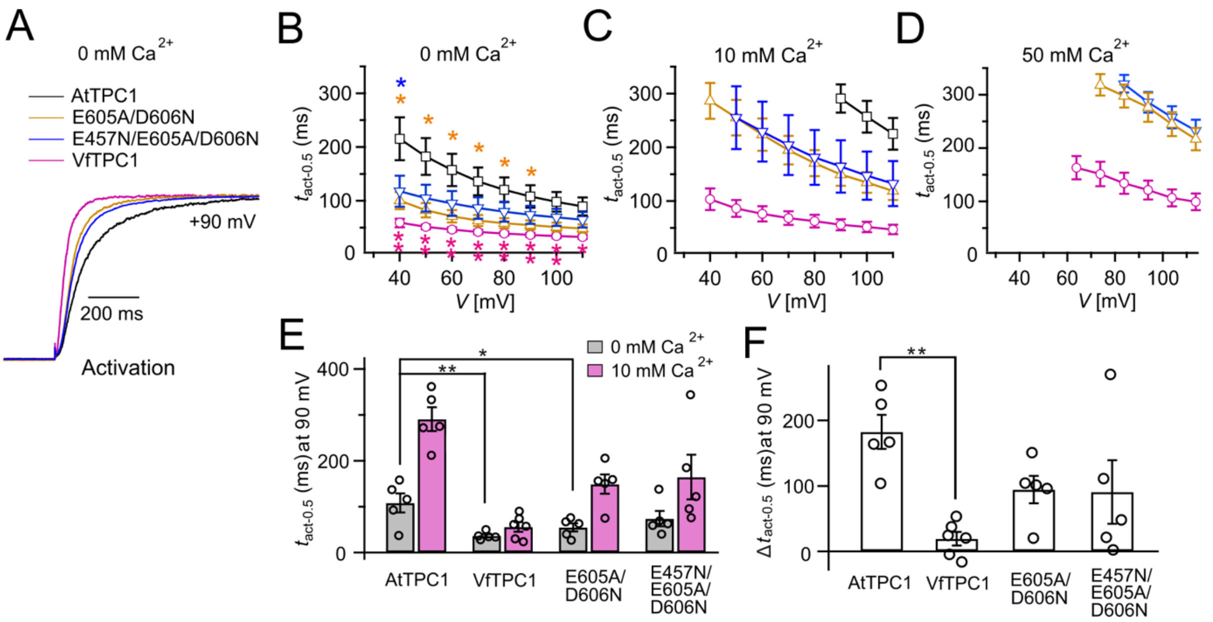


**Fig. S2.** Half-activation times of TPC1 channel variants. (*A*) Representative current relaxation induced upon a voltage pulse from -60 mV to +90 mV at 0 mM luminal $Ca^{2+}$. Normalized current responses of the vacuoles equipped with one of the indicated TPC1 channel variant were superimposed. (*B-D*) Half-activation times ($t_{act\text{-}0.5}$) of the TPC1-mediated SV currents at indicated luminal $Ca^{2+}$ concentrations plotted against the respective membrane voltages. In **b** stars denote that the $t_{act\text{-}0.5}$ values of VfTPC1 (magenta), the ATPC1 double mutant E605A/D606N (yellow) and AtTPC1 triple mutant (blue) significantly differ from AtTPC1 wild type (one-way ANOVA together with a Dunnett's post hoc comparison test; *$p < 0.05$; **$p < 0.01$). (*E*) Comparison of half-activation times determined at + 90 mV for 0 and 10 mM luminal $Ca^{2+}$. $t_{act\text{-}0.5}$ values under 0 mM luminal $Ca^{2+}$ were tested for significant differences with one-way ANOVA combined with a Dunnett's post hoc comparison test (*$p < 0.05$; **$p < 0.01$). (*F*) The differences in the half-activation times from *E* between 0 and 10 luminal $Ca^{2+}$ (one-way ANOVA together with a Dunnett's post hoc comparison test; **$p < 0.01$). In *B-F* the number of experiments (n) was identical with Fig.1*B*.

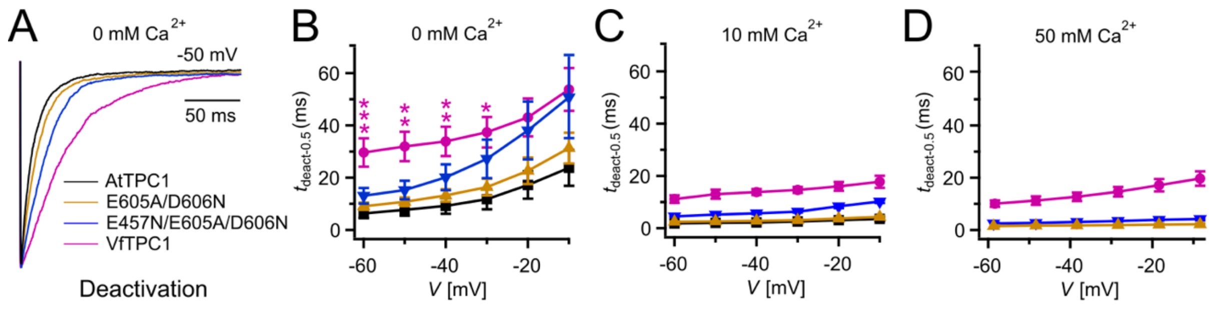


**Fig. S3.** Half-deactivation times of TPC1 channel variants. (*A*) Representative current relaxation induced upon a voltage pulse from + 80 mV or +100 mV to -50 mV at 0 mM luminal $Ca^{2+}$. Normalized current responses of the vacuoles equipped with one of the indicated TPC1 channel variant were superimposed. (*B-D*) Half-deactivation times ($t_{deact\text{-}0.5}$) of the TPC1-mediated SV currents at indicated luminal $Ca^{2+}$ concentrations plotted against the respective membrane voltages. In *B* stars denote significant differences between VfTPC1 and AtTPC1 wild type (one-way ANOVA together with a Dunnett's post hoc comparison test; *$p < 0.05$; **$p < 0.01$; ***$p < 0.001$). In *B-D* the number of experiments (n) was as follows: AtTPC1 wild type $n_{0Ca}$ = 5, $n_{10Ca}$ = 3; VfTPC1 wild type $n_{0/10Ca}$ = 4, $n_{50Ca}$ = 6; AtTPC1-E605A/D606N $n_{0\ /50Ca}$ = 4, $n_{10Ca}$ = 5; AtTPC1-E457N/E605A/D606N $n_{0/10Ca}$ = 5, $n_{50Ca}$ = 4.

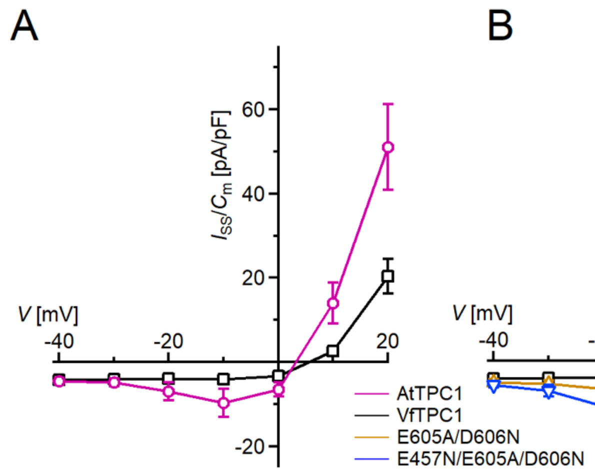


**Fig. S4.** Species-dependent effect on voltage activation threshold of TPC1/SV currents. (*A*) Enlarged section of the current-voltage plots ($I_{ss}/C_m$(V)) for AtTPC1 wild type and VfTPC1 shown in Fig. 1*B* at 0 mM luminal $Ca^{2+}$. Symbols represent means ± SE. Squares = AtTPC1 wild type with n = 5; circles = VfTPC1 wild type with n = 5. (*B*) Enlarged section of the current-voltage plots ($I_{ss}/C_m$(V)) for AtTPC1 wild type and mutant channels shown in Fig. 1*B* at 0 mM luminal $Ca^{2+}$. Symbols represent means ± SE. Squares = AtTPC1 wild type; upright triangles = AtTPC1-E605A/D606N with n = 5; reversed triangles = AtTPC1-E457N/E605A/D606N Number of experiments with individual vacuoles for each channel type was n = 5. AtTPC1 wild-type data at 0 mM $Ca^{2+}$ are identical to those shown in (1).

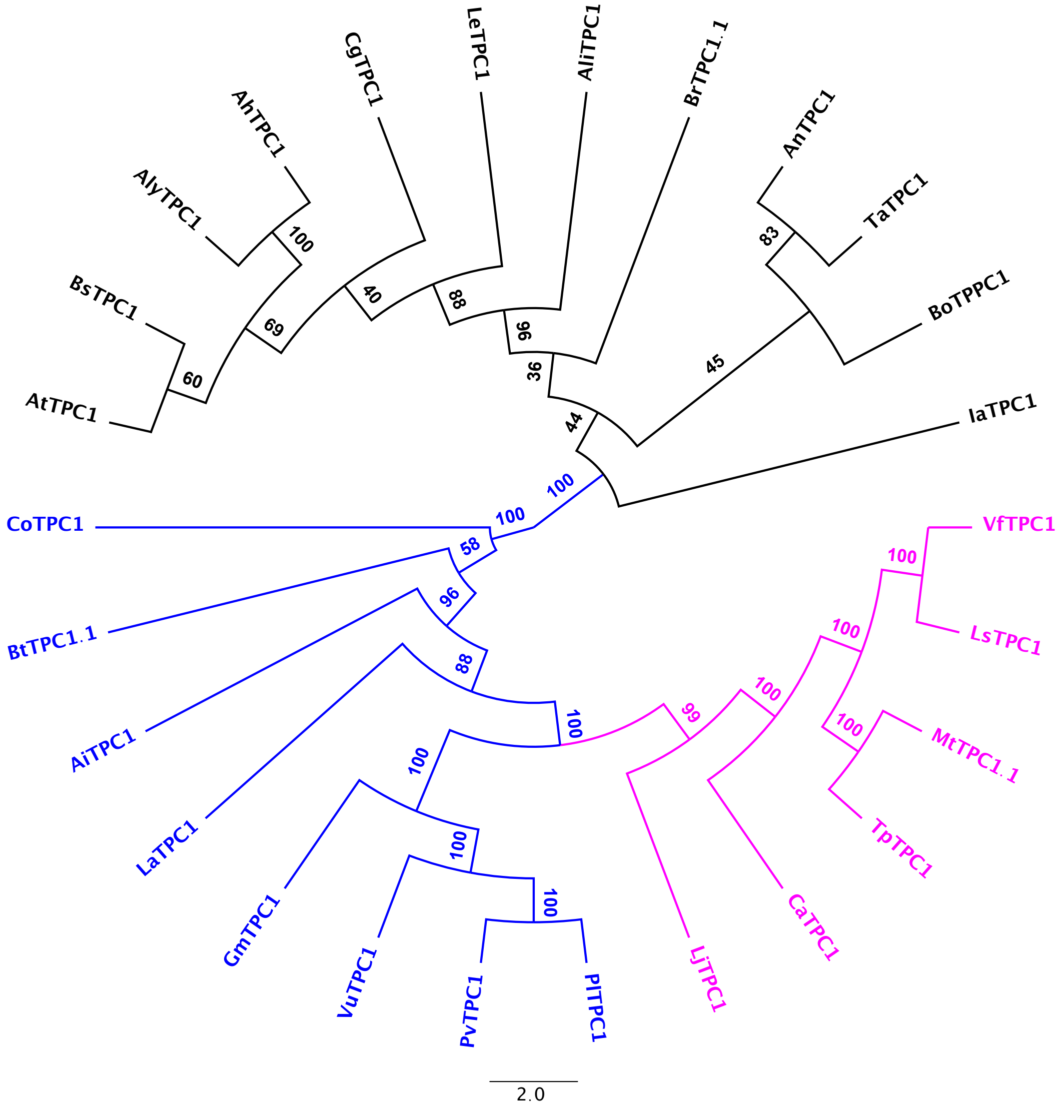


**Fig. S5.** Consenus tree of Brassicacea and Fabacea TPC1 channel proteins. Brassicacea TPC1 channels are given in black. Fabacea TPC1 channels divide into two subgroups, highlighted in magenta and blue. After a protein alignment derived with Muscle, an IQ-tree (http://iqtree.cibiv.univie.ac.at) was generated with iTol (vers. 5.7; https://itol.embl.de). Amino acid sequences and their sources are listed in File S2.

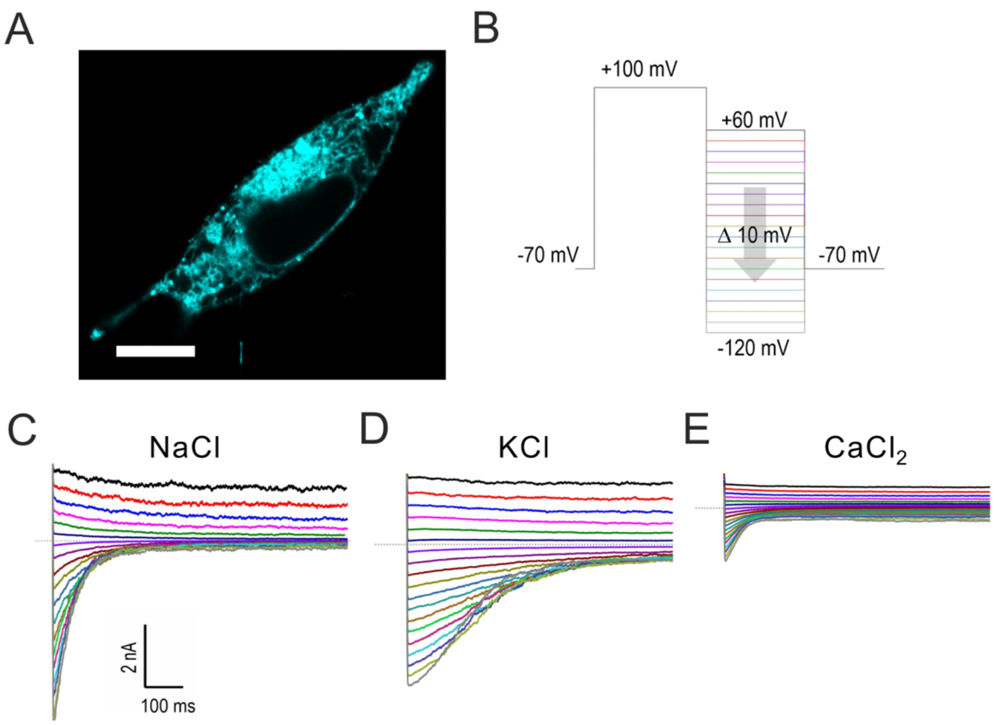


**Fig. S6.** Characterization of VfTPC1 expressed in HEK293 cells. (*A*) CLSM image of a HEK293 cell expressing eYFP-VfTPC1. Note, that most fluorescence is observed in intracellular membranes but is also present in the plasma membrane. (*B*-*E*) Electrophysiologcial characterization of VfTPC1 in HEK293 cells by whole-cell patch-clamp techniques. (*B*) Schematic overview of the protocol used during analysis including an initial activation step (-70 mV to +100 mV) and a subsequent voltage gradient (+60 mV to -120 mV) for determining the voltage dependency of TPC1. (*C*-*E*) Typical current traces of the same cell in different extracellular solutions as indicated.

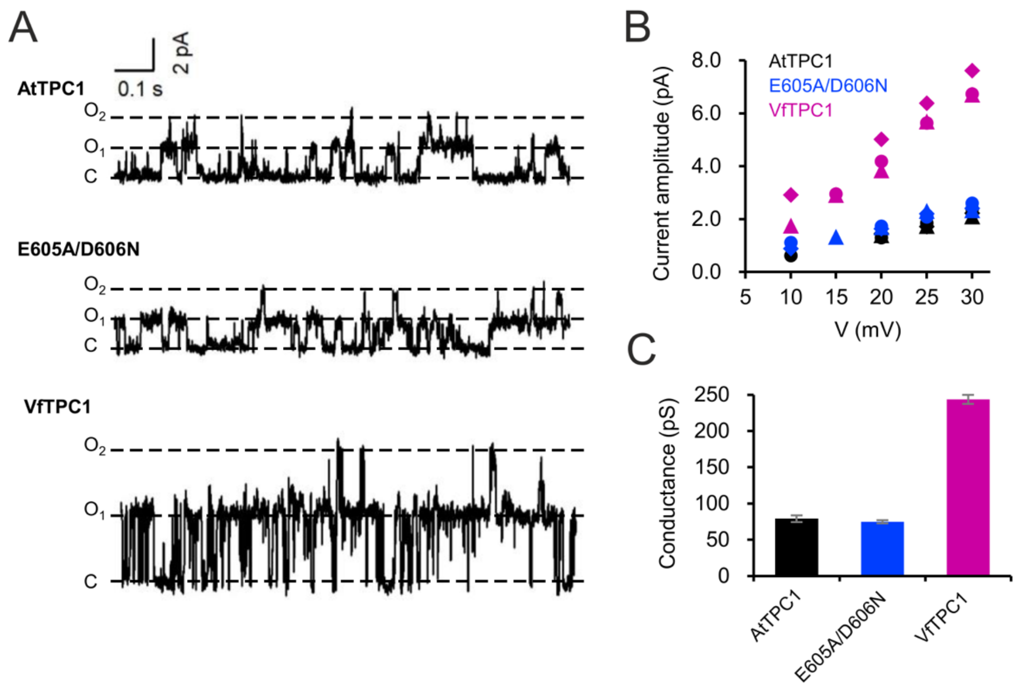


**Fig. S7.** Single channel analysis of TPC1 channel variants. (*A*) Single channel currents of indicated TPC1 channel variants evoked at +25 mV. The current level C indicates at which all channels were closed, while at current level $O_1$ and $O_2$ one or two TPC1 channels were open. (*B*) Single channel current amplitudes determined for indicated TPC1 channel variants plotted against the respective voltages. The channel variants are shown in different color codes. Each different symbol of the same colour corresponds to an individual experiment. (*C*) Bar diagram (means ± SE) gives the unitary conductance of different TPC1 channel variants derived from the linear regression fit of the individual experiments. The number of experiments was n = 3 for each channel variant.

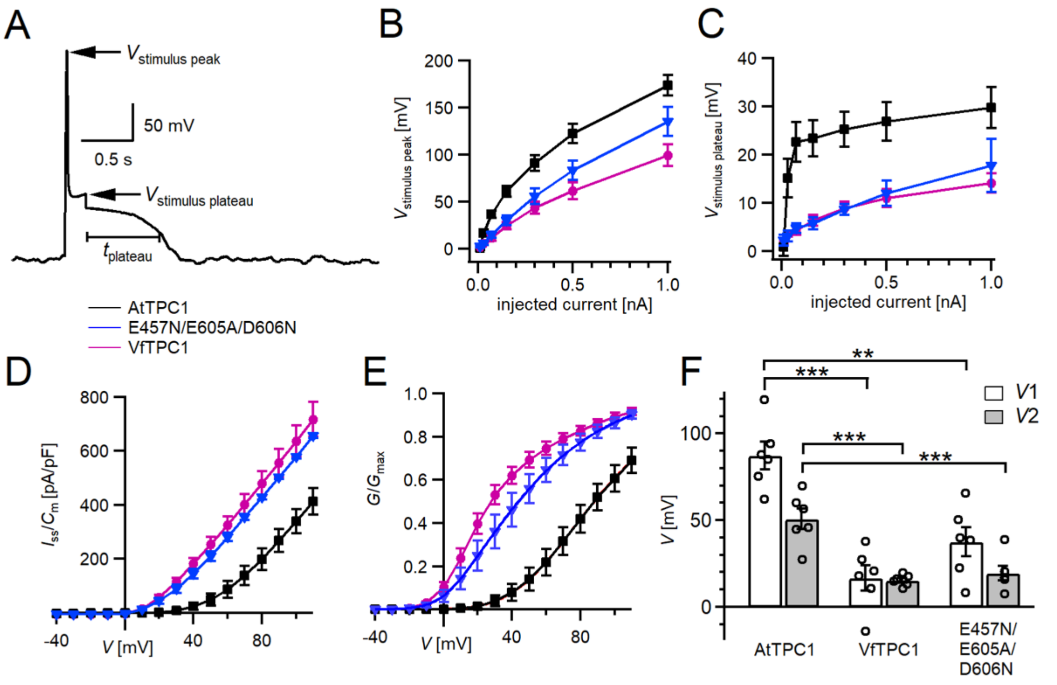


**Fig. S8.** Excitability of vacuoles with corresponding TPC1 channel activity. (*A*) Representative voltage recording from an AtTPC1-wild-type vacuole before, during and after 100-pA current injection. The voltage peak and plateau levels during current stimulation ($V_{stimulus\ peak}$, $V_{stimulus\ plateau}$, respectively) and the duration of the post-stimulus depolarization phase ($t_{plateau}$) are indicated. (*B* and *C*) Amplitudes of peak (*B*) and plateau (*C*) voltages during current stimulation recorded from vacuoles equipped with the indicated TPC1 channel variants. (*D*) Normalized currents ($I_{ss}/C_m$) mediated by the indicated TPC1 channel variant plotted against the respective membrane voltages. (*E*) Normalized conductance-voltage plots ($G/G_{max}$ (*V*)) determined for the different TPC1 channel variants as a measure for their relative open-channel probability. Best fits of the *G*/*V* plots to a double Boltzmann function are given by the solid lines. (*F*) Midpoint voltages $V_1$ and $V_2$ derived from the fits of the G(V) curves shown in *E*. $V_{1/2}$ values were tested for significant differences with one-way ANOVA followed by a Dunnett's post hoc comparison test (***p* < 0.01, ****p* < 0.001). In *B-F* symbols represent means ± SE (squares = AtTPC1 wild type; circles = VfTPC1 wild type; triangles = AtTPC1-E457/E605A/D606N). Number of experiments with individual vacuoles for each channel type was n = 6. All experiments were carried out at 0.2 mM luminal $Ca^{2+}$.

| Ca$^{2+}$ sensor sites | 2 1 2 | 1 2 | 1 | 3 3 3 |
|---|---|---|---|---|
| AtTPC1 | 239 E D | 454 D I E E | 528 E | 605 E D D |
| OsTPC1 | 264 E D | 479 D I E N | 553 E | 630 S N D |
| HvTPC1 | 252 E D | 466 D I E N | 540 E | 617 S N D |
| ZmTPC1 | 257 E D | 472 D I E N | 546 E | 623 G N D |
| SbTPC1 | 259 E D | 474 D V E N | 548 E | 625 G N D |
| | * * | * : . . | * | . . * |

**Fig. S9.** Amino acid sequence comparison of TPC1 channels in the region of the three Ca$^{2+}$ sensor sites. The numbers above the residues of AtTPC1 indicate to which Ca$^{2+}$ sensor site they contribute. A different color code was used for different amino acids. An asterisk marks 100% conserved residues across the sequences while a colon (:) indicates a conservative and a dot (.) denotes a non-conservative substitution. TPC1 sequences from following species were used: *Arabidopsis thaliana* (AtTPC1), *Oryza sativa* (OsTPC1), *Hordeum vulgaris* (HvTPC1), *Sorghum bicolor* (SbTPC1), *Zea mays* (ZmTPC1). Amino acid sequences and their sources are listed in File S2.

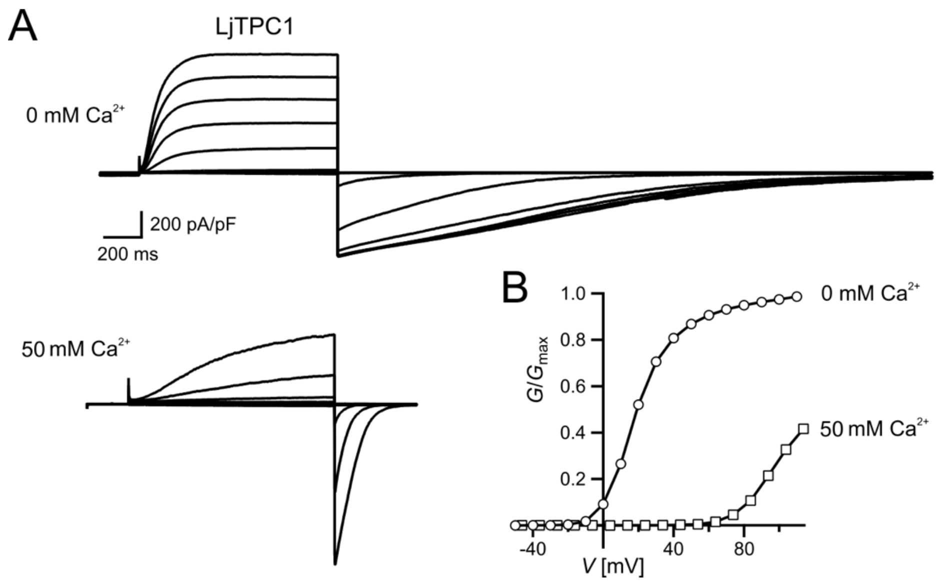


**Fig. S10.** Similar voltage-dependent gating behavior of TPC1 from *Vicia faba* and *Lotus japonicus*. (*A*) Macroscopic TPC1 current recordings from mesophyll vacuoles after transient transformation of *Arabidopsis thaliana* mesophyll protoplasts (*attpc1-2*) with LjTPC1 from *Lotus japonicus*. TPC1 currents elicited upon depolarizing voltages pulses in the range -80 mV to +110 mV in 20 mV increments. (*B*) Normalized conductance-voltage plots ($G/G_{max}$ ($V$)) determined for LjTPC1 in the absence and presence of luminal $Ca^{2+}$. Best fits of the $G/V$ plots to a double Boltzmann function are given by the solid lines.

.

**Table S1.** TPC1-dependent lifetime of the post-stimulus depolarization phase

| **Current stimulus (pA)** | **VfTPC1 $t_{plateau}$ (s)** | | **AtTPC1-triple mutant E457/E605A/D606N $t_{plateau}$ (s)** | | | | |
|---|---|---|---|---|---|---|---|
| | Exp. 1-6 | mean ± SEM (n = 6) | Exp. 1-3 | Exp. 4 | Exp. 5 | Exp. 6 | mean ± SEM (n = 6) |
| 10 | 10.00 | **10.00 ± 0** | 10.00 | 0.28 | 0.48 | 0.04 | **5.13 ± 2.18** |
| 30 | 10.00 | **10.00 ± 0** | 10.00 | 10.00 | 0.59 | 0.04 | **6.77 ± 2.04** |
| 70 | 10.00 | **10.00 ± 0** | 10.00 | 10.00 | 10.00 | 0.09 | **8.35 ± 1.65** |
| 150 | 10.00 | **10.00 ± 0** | 10.00 | 10.00 | 10.00 | 0.23 | **8.37 ± 1.63** |
| 300 | 10.00 | **10.00 ± 0** | 10.00 | 10.00 | 10.00 | 0.40 | **8.40 ± 1.60** |
| 500 | 10.00 | **10.00 ± 0** | 10.00 | 10.00 | 10.00 | 2.70 | **8.78 ± 1.22** |
| 1000 | 10.00 | **10.00 ± 0** | 10.00 | 10.00 | 10.00 | 10.00 | **10.00 ± 0.00** |
| **Current stimulus (pA)** | **AtTPC1 wild type $t_{plateau}$ (s)** | | | | | | |
| | Exp. 1 | Exp. 2 | Exp. 3 | Exp. 4 | Exp. 5 | Exp. 6 | mean ± SEM (n = 6) |
| 10 | 0.05 | 0.05 | 0.06 | 0.02 | 0.12 | 0.05 | **0.06 ± 0.13** |
| 30 | 0.08 | 0.06 | 0.12 | 0.04 | 0.25 | 0.09 | **0.11 ± 0.03** |
| 70 | 0.10 | 0.07 | 0.15 | 0.05 | 0.26 | 0.12 | **0.13 ± 0.03** |
| 150 | 0.14 | 0.08 | 0.22 | 0.05 | 0.28 | 0.21 | **0.16 ± 0.04** |
| 300 | 0.19 | 0.08 | 0.31 | 0.06 | 0.36 | 0.28 | **0.21 ± 0.05** |
| 500 | 0.38 | 0.08 | 0.40 | 0.06 | 0.48 | 0.45 | **0.31 ± 0.08** |
| 1000 | 0.55 | 0.09 | 0.53 | 0.09 | 0.62 | 0.68 | **0.42 ± 0.11** |

**Table S2.** Cation permeability of TPC1 channel variants

| Channel variant | $P_K/P_{Na}$ Mean ± SD (n) | $P_{Ca}/P_{Na}$ Mean ± SD (n) |
|---|---|---|
| AtTPC1 wild type | 0.75 ± 0.08 (5)*** | -- |
| AtTPC1$^{E605A/D606A}$ | 0.85 ± 0.14 (6)** | 5.43 ± 0.69 (4) |
| AtTPC1$^{D240A/D454A/E528A}$ | 0.85 ± 0.09 (5)** | 5.50 ± 0.45 (4) |
| VfTPC1 wild type | 1.18 ± 0.11 (4) | 5.92 ± 0.52 (4) |

Significant differences of $P_K/P_{Na}$ between AtTPC1 channel variants and VfTPC1 (**$p$ < 0.01; ***$p$ < 0.001) determined with one-way ANOVA together with a Dunnett's post hoc comparison test.

**Table S3.** Primer sequences for subcloning and side-directed mutagenesis

| Name | Sequence (5'- 3') |
|---|---|
| AtTPC1$^{WT}$ user fwd | GGCTTAAUATGGAAGACCCGTTGATTGG |
| AtTPC1$^{WT}$ user rev | GGTTTAAUTCATGTGTCAGAAGTGGAACACTC |
| AtTPC1$^{D240A}$ fwd | AGGCCACUCAGCAGGGCCTCACGGTC |
| AtTPC1$^{D240A}$ rev | AGTGGCCUCAAACATAACAAAAGCAATCCAA |
| AtTPC1$^{D454A}$ fwd | ACGCTTGCTAUCGAAGAAAGCTCGGCTCAG |
| AtTPC1$^{D454A}$ rev | ATAGCAAGCGUTGTTTCAACAACGACAGCAA |
| AtTPC1$^{E457N}$ fwd | AAACAGCUCGGCTCAGAAGCCATGG |
| AtTPC1$^{E457N}$ rev | AGCTGTTUTCGATATCAAGCGTTGTTTCAA C |
| AtTPC1$^{E528A}$ fwd | AGCATGGAUCCGGTACCTTCTCCTGGC |
| AtTPC1$^{E528A}$ rev | ATCCATGCUCCATTTGAGAAGAAAGTATTCTCG |
| AtTPC1$^{E605A}$ fwd | ATTGGCTGCGGAUGACTACCTTTTGTTCAAC |
| AtTPC1$^{E605A}$ rev | ATCCGCAGCCAAUTCGGTTTCAAA |
| AtTPC1$^{D606N}$ fwd | AGAGAATGACUACCTTTTGTTCAACTTCAAT |
| AtTPC1$^{D606N}$ rev | AGTCATTCTCUGCCAATTCGGTTTCAAAGAG |
| VfTPC1$^{WT}$ user fwd | GGCTTAAUATGACGGAGCCTCTACTCAGAG |
| VfTPC1$^{WT}$ user rev | GGTTTAAUCCTGTACTGGAAGGATGATTTTGACAAA |
| LjTPC1$^{WT}$ user fwd | GGCTTAAUATGGAACCTCTGCTCAGAGGCGAAAGCAGTG |
| LjTPC1$^{WT}$ user rev | GGTTTAAUTTATGCATTGGAAGGCTGATCTTGACACAACTCAG |

fwd = forward primer, rev = reversed primer
